# Dihydroartemisinin induces a two-step transcriptional response and stage-specific developmental shifts in malaria parasites, *Plasmodium falciparum*

**DOI:** 10.64898/2026.06.16.732544

**Authors:** Josephine Boentoro, Michal Kucharski, Sourav Nayak, Zbynek Bozdech

**Affiliations:** School of Biological Sciences, Nanyang Technological University, Singapore; Mahidol-Oxford Research Unit, Bangkok, Thailand; Department of Clinical Tropical Medicine, Mahidol University, Thailand

## Abstract

The parasite *Plasmodium falciparum* causes malaria, the deadliest human parasitic disease, which remains fatal when not promptly treated. Evolving parasite resistance to frontline artemisinin-based therapies threatens vulnerable populations and decades of progress toward malaria elimination. Yet the mode of action of dihydroartemisinin (DHA), the active metabolite of these treatments, remains incompletely understood. Here we applied dose-response transcriptomics across the three intraerythrocytic stages - rings, trophozoites, and schizonts, revealing a two-tier transcriptional response to DHA, with low- and high-dose programs consistent with specific drug action and cytotoxic damage. The trophozoite stage mounts the strongest and most coordinated response, including a striking reversal of the developmental cascade in which protein synthesis machinery is broadly downregulated and a ring-like transcriptional profile is reactivated - reminiscent of drug-induced quiescence. Coordinated regulation of multiple protein complexes, most notably Kelch13 and its interacting partners (KIC), points to organized transcriptional control of the parasite’s drug response. This work provides a stage- and dose-resolved view of DHA action in *P. falciparum* and a template for future antimalarial mechanism-of-action studies.

## Introduction

Malaria is a severe disease caused by infection with *Plasmodium* parasites, most commonly *P. falciparum and P. vivax.* In 2024, there were an estimated 282 million malaria cases across 80 endemic countries, resulting in 610 thousand fatalities, representing an estimated increase of 9 million cases from 2023^1^. The frontline antimalarial treatment currently implemented in all malaria-endemic regions worldwide is artemisinin combination therapy (ACT), which typically consists of a fast-acting artemisinin derivative paired with a slow-acting partner drug, typically based on quinoline scaffolds^2^. The active metabolite of artemisinin, dihydroartemisinin (DHA), is highly potent against parasites throughout the *Plasmodium* intraerythrocytic development cycle (IDC) of *Plasmodium falciparum*, having a half-maximal inhibitory concentration (IC50) of below 10 nM^3^. Artemisinin and its derivatives are believed to exert their parasiticidal effects when the endoperoxide bridge is activated by heme, producing free radicals. These subsequently cause mitochondrial membrane depolarization and widespread damage to proteins and lipids, leading to cellular death^4^. Artemisinin compounds are effective across all stages of the IDC, albeit with varying IC50 values, likely reflecting differences in the blueprint of the parasite’s cellular physiology at different stages of asexual intraerythrocytic development^5–7^.

Despite its effectiveness, partial resistance of *P. falciparum*, the deadliest species of malaria parasites, to artemisinins was reported in Southeast Asia as early as 2009^8^ and has likely evolved independently in sub-Saharan Africa about a decade later^9,10^. In clinical settings, parasites are considered resistant when exhibiting a clearance half-life ≥5h hours^11^. *In vitro*, resistance is measured using the ring-stage survival assay, which quantifies the number of parasites that survive a 6-hour pulse of 700 nM DHA in very young (0-3 hours post-invasion, HPI) rings^12^. This resilience of early ring stage parasites to artemisinin (e.g., artemisinin partial resistance) is believed to underlie the ultimate rise of ACT treatment failures, by selecting resistance alleles to the partner drugs^13^. Due to this development, understanding the molecular mechanism by which the resistant parasite withstands the putative polypharmacological mode of action (MOA) of artemisinin is of great importance for the better management of current ACTs that remain efficacious for the time being ^14^. Such knowledge will also be important for the design of alternative ACT formulations, such as Triple ACTs (TACT), which will be urgently needed to protect the utility of artemisinin-based chemotherapies^15^. This will be particularly important given that no new antimalarial chemotherapeutic agent is expected to reach full implementation for the upcoming years.

The mechanism of artemisinin resistance has been a subject of ongoing research, with the strongest association with nonsynonymous mutations in *pfkelch13*^16–18^. These mutations have been associated with changes in hemoglobin endocytosis^19^ and reduced phosphatidylinositol-3-kinase (PI3K) ubiquitination^20^. The corresponding increase in phosphatidylinositol-3-phosphate (PI3P) has also been shown to increase tolerance to artemisinin *in vitro*. Parasites bearing *pfkelch13* mutations introduced by gene editing also exhibit differential cellular states in the absence of an artemisinin-resistant (ART-R) genetic background. These include increased protein turnover, altered carbon metabolism, and a prolonged ring stage with the C580Y mutation^21^. However, it is becoming progressively clear that *pfkelch13* polymorphisms are not the sole determinants of artemisinin resistance. In a landmark study of ART-R clinical isolates, Miotto *et al.* demonstrated that nonsynonymous *pfkelch13* mutations tended to occur on a distinct genetic background^17^. The effectiveness of nonsynonymous *pfkelch13*mutations also varies depending on the parasite strain and, hence, the genetic background on which they are tested^18^. This concept was further expanded upon by the association of eQTLs and variations in noncoding regions with artemisinin resistance^22–24^. In addition, various studies have identified other genetic variations that confer increased tolerance to artemisinin independently of *pfkelch13*, such as coronin^25^, falcipain-2a^26^, ubp1^27^, ap2mu^28^, and others^29^. In spite of these discoveries, a comprehensive understanding of the molecular mechanisms that mediate artemisinin resistance and its polypharmacological MOA remains a major knowledge gap for the better management and implementation of current and future ACTs.

In contrast to standard model systems, the utility of transcriptomics for studies of *Plasmodium* parasites gained traction only slowly through the first decade of the 21^st^ century^30^. Indeed, it was initially thought that malaria parasites, particularly *P. falciparum*, possess a “hard-wired RNA metabolism” when treatment with external perturbations, such as a DHFR inhibitor (pyrimethamine), failed to induce a significant transcriptomic response^31^. Furthermore, the few genes that were observed to be differentially expressed appeared unrelated to the compound’s known target and were termed the “RNA whispers”. This limited transcriptional response has since been understood to be a function of the nature of the perturbator and not generalizable to *P. falciparum* biology^30,32^. Since then, improvements in transcriptomic technologies and the increased availability of RNA-Seq have enabled high-throughput detection and absolute quantification of transcripts, supplanting earlier microarray and qPCR techniques^32,33^. Subsequently, many studies have been published on the *P. falciparum* transcriptional response to numerous external pressures, including drugs and chemical agents, showing that while in some cases *P. falciparum* indeed does not respond, in others highly informative transcriptional changes are directly linked to the perturbation MOA^30^. Given the high importance of artemisinin, a sizable number of previous studies investigated transcriptional responses of *P. falciparum* to artemisinin *in vitro*^21,29,34–39^ and to a smaller extent *in vivo*^22,24^. Indeed, several key biological processes, such as protein synthesis and turnover, redox homeostasis and energy metabolism, mitochondrial function, etc., were found to be transcriptionally dysregulated due to artemisinin exposure. In all these studies, however, transcriptional responses were induced by a single-dose exposure of DHA, set at either a low, near-50% inhibitory concentration (IC50) of 5-50nM or a physiologically relevant 700nM, typically focusing on one or a limited number of IDC development stages (ring, trophozoite and schizont) at a time. Due to this, each study captured a certain aspect of artemisinin’s parasiticidal bioactivity within a certain window of the IDC. Although each of these studies provided valuable insights, the full picture of the dose- and time-responsiveness of *P. falciparum* across the entire IDC remains incomplete. Dose responsivity is of particular interest, given that the true physiological concentration is known to vary greatly, as pharmacokinetic profiles may vary between individuals^41,43–45^.

In this study, we aimed to help close this knowledge gap by performing transcriptional profiling of *P. falciparum* across the most active derivative, DHA, in a dose-dependent manner over several time points at three IDC development stages: rings, trophozoites, and schizonts. To investigate the concentration effect on the drugs’ MOA, we utilized the Dose-Response OMICS (DROMIC) approach^40,41^ and demonstrate that the *P. falciparum* transcriptional responses to DHA are highly dose-responsive, and this responsiveness varies dramatically across the IDC. By integrating temporal analysis, the DROMIC approach also allowed us to distinguish function-specific transcriptional responses from generic shifts in IDC transcriptional cascades, both of which are induced by DHA in *P. falciparum* parasites.

## Results

### Profile of the DHA-induced transcriptome

The key rationale for this experimental design was to provide a comprehensive picture of the transcriptional responses of *P. falciparum* to the most active artemisinin derivative (DHA) in both dose- and time-dependent manners at three IDC stages. The entire study presented here is based on a single experimental approach in which *an in vitro*-cultured *P. falciparum* 3D7 strain was treated with increasing concentrations of DHA ranging from 1.9 nM to 150 in threefold serial dilutions. These dose-response treatments were conducted at the three main IDC stages: ring (8 hours post invasion, HPI), trophozoite (16 HPI), and schizont (36 HPI) (**Figure 1A**, & **Materials and Methods).** For the transcriptomic analysis, cell samples were harvested at the beginning of the experiment (0 hrs) and at 1, 4, and 6 hrs after treatment at each DHA concentration. For the overall RNA-Seq analysis, RNA was isolated from a total of 171 samples (3 biological replicates, 3 IDC stages, 5 treatment concentrations + 1 untreated control, 3 treatment durations + 1 initial sample). The resulting RNA samples were processed for massively parallel sequencing on the Illumina Novaseq 6000 platform, using Smart-Seq2 amplification and the Illumina Nextera XT library preparation kit, with previously optimized methods^42^. After mapping and quantification of the resulting raw read count datasets from the RNA-Seq analysis, the transcript per million (TPM) and raw read count values were obtained for individual mRNA transcripts. After low-expression filtering of protein-coding transcripts (**Materials and Methods**), we obtained mRNA abundance profiles of 4153, 5207, and 5378 transcripts in rings, trophozoites, and schizonts, respectively (**Figure S1-3**). To identify genes that respond to the DHA treatments in a dose-responsive manner, the transcriptional profiles within the concentration gradient were first analyzed by the DRomics package^43^, evaluating dose-response transcriptional profiles at each time and IDC experimental point separately (**Figure 1A**). Transcripts were first tested for a quadratic relationship between dose and RNA-seq read counts with an FDR cutoff of 0.05. Dose-responsive transcripts were then fitted to five models: linear, exponential, Hill, Gauss-probit, and log-Gauss-probit (**Figure S4 & 5**). mRNA transcripts whose abundance profile exhibited a fit to at least one of these models were classified as dose-responsive and thus included in the following functional analyses. Moreover, we also used the transcritonal profiles of the initial untreated parasite control samples (H0) to confirm the correct IDC stage by correlation with a reference IDC transcriptional cascade **(Figure S6)**.

**Figure 1.**
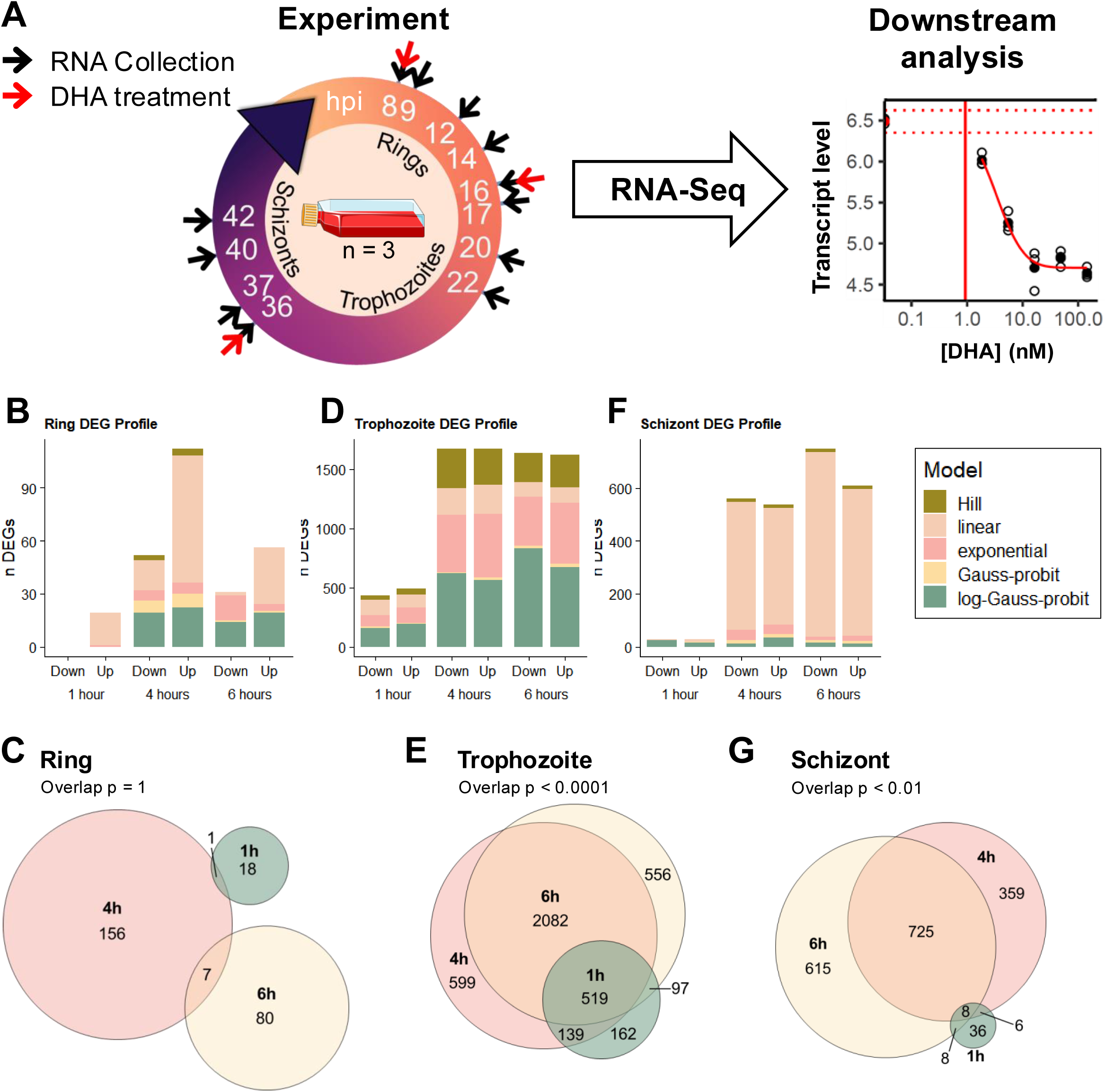
Profile of the DHA-responsive transcriptome. Description: A) Experiment design: a range of 0-150 nM DHA is added to synchronized parasites at 8, 16 and 36 hours post-invasion (red arrows). RNA is harvested at 0, 1, 4, and 6 hours post-treatment (black arrows) for RNA-Seq. Transcripts are fitted to curve models and the point of departure calculated in downstream analysis. B-D) Bar graphs indicating the total number of differentially expressed genes and the best-fit model (color) for each treatment duration and in rings (B), trophozoites (C), and schizonts (D). E-F) Proportional Venn diagrams of differentially-expressed genes between each treatment duration per stage. P-values were calculated against a multivariate hypergeometric distribution for the overlap of three sets of genes.

Overall, we observed stark differences in the extent of DHA transcriptional responses among the three IDC stages, with the greatest number of differentially expressed genes (DEGs) in trophozoites, followed by schizonts and rings (**Figure 1B-F**). At the ring stage, there were 19 upregulated and zero downregulated genes after 1 hr of DHA treatment, suggesting a near complete absence of an immediate transcriptional response (**Figure 1B**). There was a considerable increase, however, in the number of DEGs at the 4-hr treatment course, with 112 and 52 up- and down-regulated genes, respectively. Curiously, after 6 hrs of DHA treatment, the number of DEG dropped to 56 up- and 31 down-regulated genes. There was, however, no statistically significant overlap in the DEGs at the three exposure time points in the ring stage, calling into question the overall specificity and coordination of the ring-stratified transcriptional response to DHA (**Figure 1C**). On the other hand, in trophozoites, we observed a substantial and robust transcriptional response as early as 1 hour, where 488 and 429 genes were upregulated and downregulated, respectively (**Figure 1D**). Subsequently, we detected 1668 and 1671 up- and down-regulated genes at 4 hours of DHA treatment, and, similarly, 1619 and 1635 up- and down-regulated genes at the 6-hr DHA treatment (**Figure 1D**). The sets of DEGs at the trophozoite stage also exhibited highly statistically significant overlaps (p < 0.0001), suggesting the existence of robust transcriptional regulation mediating the drug response (**Figure 1E**). Hence, while the ring stages appear unresponsive, the trophozoites, on the other hand, seem to mount a broad yet highly coordinated response to DHA. In schizonts, there was a comparatively low immediate response, similar to that in rings, with 29 genes each up- and down-regulated at 1 hr (**Figure 1F**). After 4 hrs of treatment, however, the number of DEGs increased, with 537 genes upregulated and 561 downregulated. The trend continued after six hours of treatment, with 608 and 748 genes up and down-regulated, respectively. DEGs in schizonts also exhibited a significant degree of overlap (p < 0.01) (**Figure 1G**). Taken together, we demonstrate that *P. falciparum* has a strong capacity to respond transcriptionally to the frontline chemotherapeutic agent DHA in a dose- and time-dependent manner. This transcriptomic response appears highly coordinated in the trophozoite and schizont stages. In contrast, DHA fails to elicit any specific transcriptional response in rings, being reminiscent of the “RNA whisper” previously reported for transcriptional responses to pyrimethamine (*authors’ note*)^31^.

### DHA induces a two-step transcriptional response

In the next step of the analysis, we used the dose-dependent transcriptional profiles to determine the benchmark dose (BMD), which denotes the dose at which each transcript abundance deviates from baseline. BMD thus reflects the effective concentration of DHA that affects the transcriptional regulation of each gene, thereby sensing the cytotoxic effect of the drug^43,44^. Remarkably, inspection of the BMD distributions in the identified sets of DEGs revealed a clear overall separation into two groups: BMD between 0-10 nM (BMD_low_) and 50 to 100 nM (BMD_high_) (**Figure 2A**, **Table S1A**). Specifically, the BMD distributions at each stage and duration exhibit a distinct BMD_low_ population with a mean of between 0.6 and 4.7 nM DHA, and a BMD_high_ population with means between 44 and 68.1 nM DHA. Gaussian mixture modeling (k=2) revealed the presence of both BMD groups in transcriptional responses of rings (4 and 6 hr, **Figure 2B& C),** trophozoites (1 hr, **Figure 2D**) and schizonts (4 and 6 hrs, **Figure 2E& F**; 1 hr, **Figure S5**). On the other hand, the dual BMD distribution model failed to converge in trophozoites treated for 4 and 6 hours, in which most DEGs followed the BMD_low_ profile, with a median of 1.2 nM in both cases (**Figure S5**). The bimodal distribution could not be modeled in 1hr-treated rings, as there were too few (n=19) DEGs, largely belonging to the BMD_high_ group (Median = 73 nM). Schizonts treated for 1 hour show this bimodality to a greater extent, with the BMD_low_ group (n = 40) entirely composed of transcripts with BMD < 5 nM DHA. The BMD_high_ group at this stage (n = 18) shows a large gap from this, with a BMD range from 47.8 to 96.1 nM DHA. Taken together, the distribution of BMD values indicates two distinct groups of transcripts responding in two waves of the DHA concentration gradient. Notably, the concentrations representing BMD_low_ are related to the *in vitro* half-maximal inhibitory concentration (IC_50_) for *P. falciparum* growth, whereas BMD_high_ is more reflective of *in vivo* DHA concentrations (>100 nM). It is tempting to speculate that the observed transcriptional responses indeed reflect two sub-mechanisms of the artemisinin MOA, one of which is related to parasiticidal activity at low concentrations (IC_50_ <10nM) and the other in higher concentrations (>100nM), including the peak physiological concentration in patients’ peripheral blood (∼700nM).

**Figure 2.**
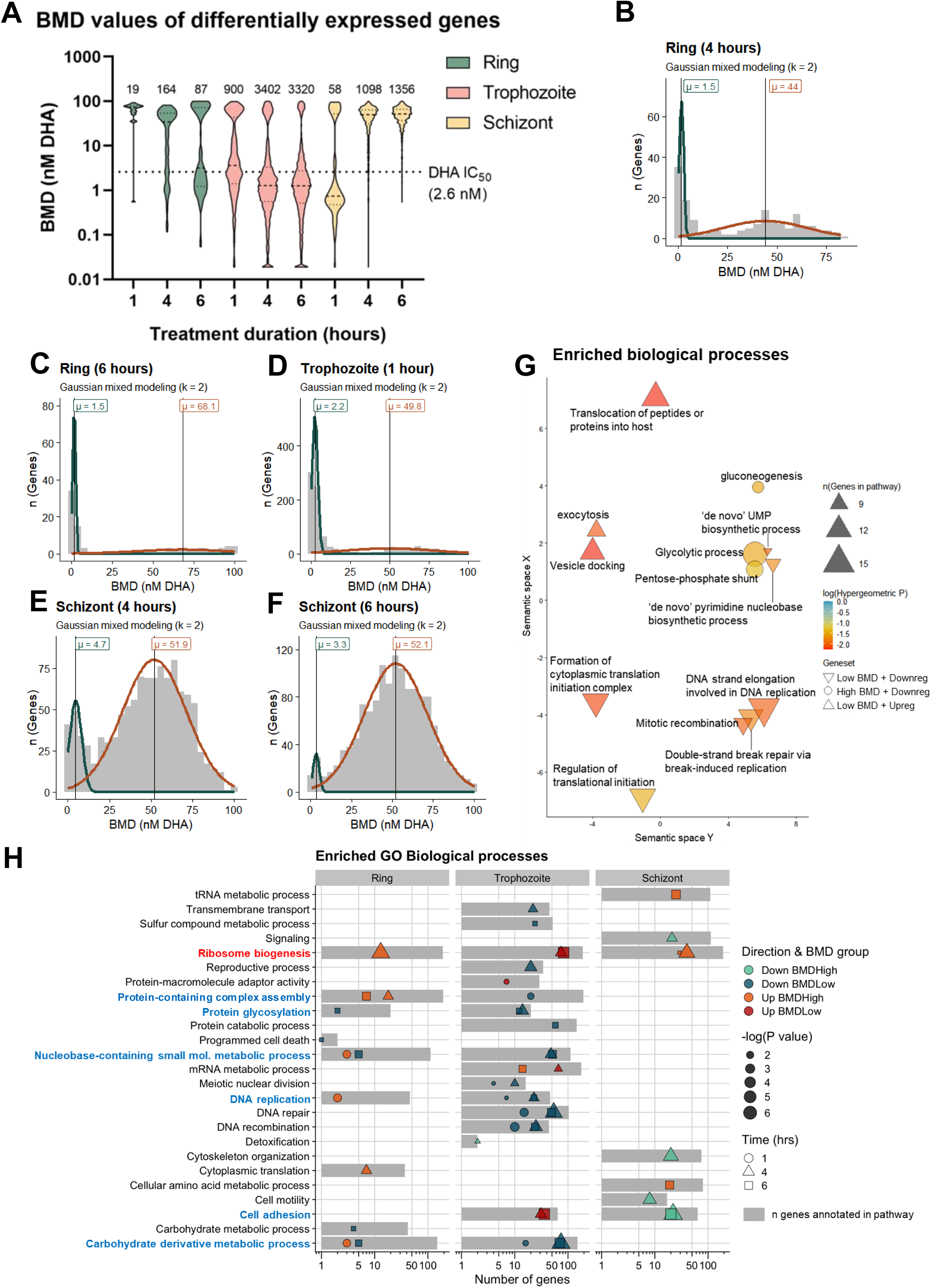
Bimodal transcriptional response to DHA. **(A)**Violin plot of the calculated BMD values (y axis) of each DE gene in each stage (color) and treatment duration (x axis), irrespective of upregulation or downregulation. The plot demonstrates a tendency for either low, near-IC_50_ BMDs, or high BMDs. Dashed lines inside violins indicate group median, with quartiles indicated by dotted lines. The IC_50_ of DHA as measured by a standard drug susceptibility assay is annotated with a dotted line across the plot. Figure generated in Graphpad Prism 8. (**B-F**) Histograms of BMD values (in nM DHA) from each stage and corresponding treatment duration that converged to a binomial model. Solid lines represent the corresponding gaussian density distribution scaled to the gene counts. The line color corresponds to either low distributions (green) or high distributions (brown). The mean of each distribution is annotated by a vertical line and text. **(G)** Semantic similarity plot of terms enriched in differentially expressed genes in trophozoites after 1 hour of treatment. Shapes indicate the direction of differential expression and the BMD group of the genes. Size of the shape indicates the number of genes annoted in the pathway, while the fill shows the adjusted hypergeometric P-value. **(H)** Summary of biological processes enriched in DE genes from three parasite stages. Gray bars indicate the number of annotated genes in the pathway. Points indicate the number of DE genes belonging to that pathway, with the colors indicating the direction of differential expression and the BMD group, size corresponding to –log(Hypergeometric P value) adjusted by Benjamini-Hochberg method, and the shape indicating the time of treatment in hours.

With the notion of the dual mode of DHA MOA, next, we subsequently carried out functional analysis of the DEGs with respect to the BMD, but also the IDC stage and time of treatment (**Table S1B**). To this end, we tested for functional enrichment of Gene Ontology (GO)- annotated pathways within each gene group (**Figure 2G-H, Tables S2-3, Supplementary Datasets 1 & 2**). First, we investigated functions enriched in trophozoites at 1 hour of DHA exposure, which represents immediate, direct transcriptional responses to the drug’s cytotoxic effect (**Figure 2G**). At BMD_low_ values, several functional groups related to vesicular flow and protein transport were upregulated, driven by upregulation of 6 SNARE complex components, 5 of which are essential^45^. On the other hand, there was a significant enrichment of factors involved in translation initiation and DNA replication amongst the genes downregulated at BMD_low_. The DHA transcriptional responses at BMD_high_ were distinct from those at BMD_low,_ with no functional overrepresentations among the upregulated genes but significant enrichments of glycolysis, gluconeogenesis, and the pentose phosphate shunt among the downregulated genes (**Figure 2G**).

In contrast to the short-term treatment, 4 and 6 hours of DHA treatments of the trophozoites induced considerably more profound changes in the downregulated genes at BMD_low,_ including factors involved in lipid metabolism, DNA replication, proteolysis, protein translation, and assembly of cellular components (**Figure 2H**). Conversely, only factors of cell adhesion, mRNA metabolism, and ribosomal biogenesis were upregulated by the 4 and 6 hr exposures of the trophozoite to DHA, the latter of which was also upregulated in the other IDC stages. The transcriptional response of rings and schizonts was much less functionally defined than trophozoites, exhibiting sparse pathway enrichments in the BMD_High_ group (**Figure 2H, Table S2**). Other processes shared between the two IDC stages are protein complex assembly, glycosylation, nucleobase metabolism, DNA replication, and cell adhesion. Taken together, the distinct classes of functional assignments amongst the individual sets of DEGs across the three IDC stages, time intervals, and BMD further reinforce the specificity and coordination of this transcriptional response as a function of DHA-induced cellular stress.

### DHA induces transcriptional cell cycle regression in trophozoites

It is well established that during the IDC, each gene of the *P. falciparum* genome is tightly regulated by being expressed at highly specific times when its function is required. As such, the overall transcriptome of the IDC represents a highly synchronized cascade-like pattern, also termed the “just-in-time” transcriptional program^46–48^. Due to the periodic nature of the basal level of essentially all mRNA profiles, all analyses of differential gene expression in *P. falciparum* must take into consideration parasite transcriptional age^49^. Indeed, DHA has been shown to affect the progression of IDC of the transcriptional cascade in *P. falciparum,* particularly at the ring and early trophozoite stages^21^. Given the comprehensiveness of the transcriptomic dataset generated in this study, we wished to explore this in greater detail. To test this, we correlated each sample with each time point from the reference IDC transcriptional cascade that reflects the dominant stage representation in each sample with nearly a single-hour resolution (**Figure 3A**, **Figure S6 & S7**). As expected, the 0-hour samples prior to treatment of the rings, trophozoites, and schizonts control samples exhibited the highest correlation with the anticipated IDC ages, corresponding to approximately 8, 23, and 33 HPI (**Figure 3A**). In the treatment courses, the ring and schizont samples at 1, 4, and 6 hours showed identical correlation profiles to those in the control samples. This indicates that in both stages, DHA has no discernible effect on IDC progression, as peak correlation values between untreated and treated samples remain at the same timepoints (**Figure 3A**, **Figure S7**). On the other hand, 4 and 6 hours of DHA exposure of trophozoites resulted in a significant increase in correlation with earlier IDC stages (10-20 hpi). This correlation profile is consistent not only with the stall in their progression during the IDC but also with a possible reversal of at least a portion of the transcriptional cascade back towards the ring stages (**Figure 3A**). This trend is consistent with the downregulation of multiple pathways related to RNA and protein metabolism and DNA replication (**Figure 2H**) whose transcriptional activity is expected to rise through the trophozoite stages under normal growth conditions^46^.

**Figure 3.**
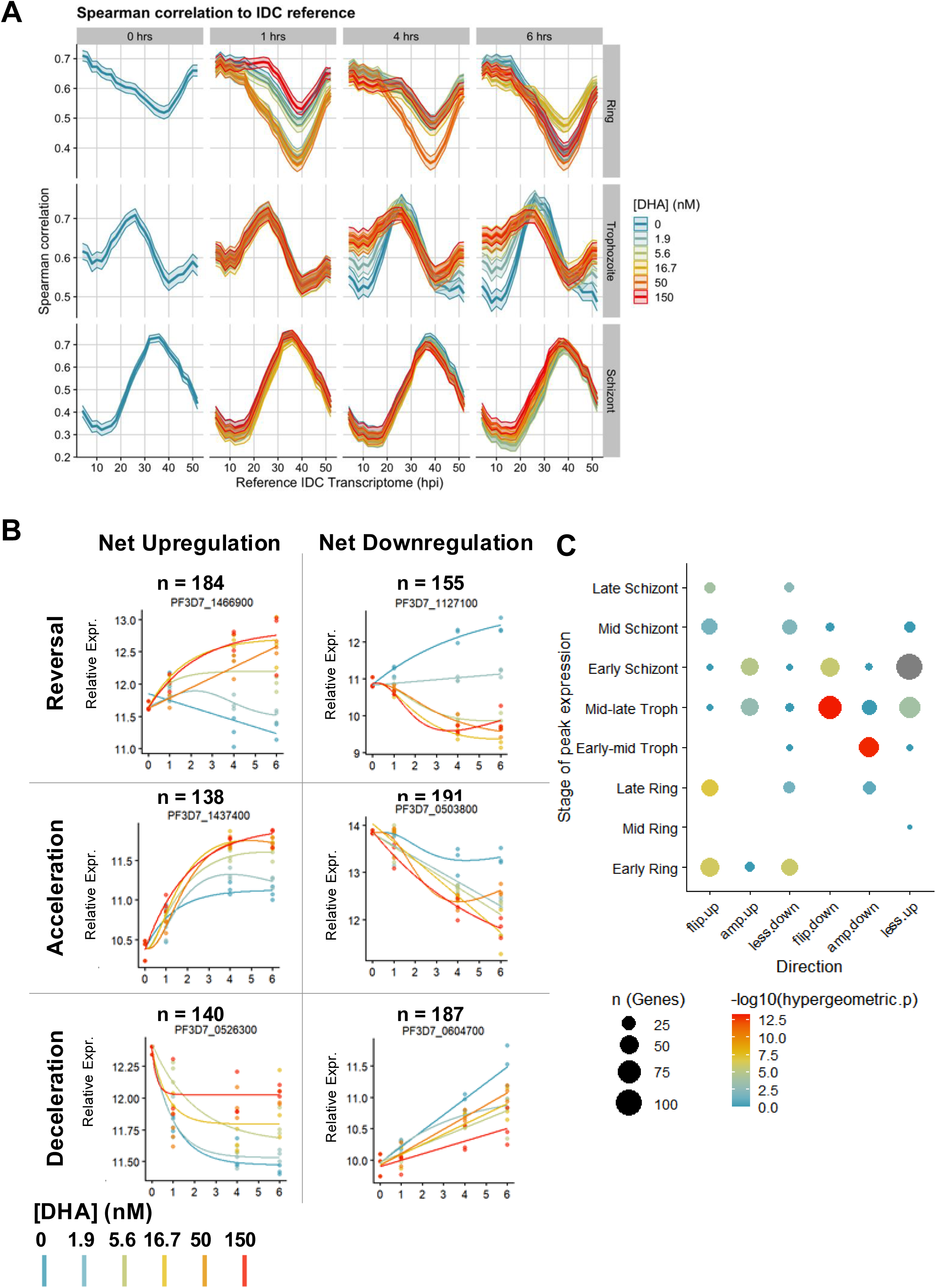
Reversal of life cycle transcriptional patterns. **(A)** Spearman rank correlation values (solid line) between experiment samples at each stage (rows) and treament duration (columns) with each sample from the reference IDC transcriptome generated by Kucharski *et al.* (2020). The x-axis indicates the hours post-invasion (hpi) of the reference samples, while ribbons around the lines indicate the 95% confidence interval derived from bootstrapping (n = 1000). Colors indicate the DHA dose of the corresponding line. **(B)** Examples of each of six dynamic differential expression patterns described in this study, demonstrating the relationship of the slope of expression over time in treated versus untreated sample. Points denote normalized expression values from three experimental replicates after VST transformation. As time (X-axis) progresses, changes in expression (Y-axis) are measured for each dose (colors). The number of genes classified per DDE pattern is denoted above each example graph by n. **(C)** Genes in each DDEP (x-axis) were tested for enrichment of genes that peak in each stage (y-axis). Values with adjusted hypergeometric *p* value (Benjamini-Hochberg) are indicated with fill color. Point size corresponds to the number of genes belonging to the stage in that DDE pattern. Red arrows below the x-axis indicate the vector (direction and relative amplitude) of gene expression after treatment relative to the untreated control (blue arrows).

Next, we hypothesized that the IDC reversal from trophozoite to rings may be achieved not only by a failure of transcript-level decline but possibly also by reinitiation of at least some ring-specific transcripts. To inspect this more closely, we reanalyze the generated transcriptomes using the DRomics pipeline, with time as the primary variable (for details, see **Materials and Methods**). Here we modeled expression changes for each transcript across each dose across the 0, 1, 4 and 6 hr time course . Comparing the simple 6-hour slope of a transcript at high DHA doses to that in the corresponding untreated samples enabled classification into six dynamic differential expression patterns (DDEPs) (**Figure 3C**). These include two categories with reversed expression patterns: “flip up” -genes with declining profiles through the trophozoite stage are reinitiated across the 6 hr treatment course (n=184); and conversely, “flip down” -genes suppressed by DHA treatments despite their increasing endogenous pattern in untreated parasites (n=155) (**Figure 3B**, top row). In addition, we classified four DDEPs with altered but not reversed trend, including “amp up” -genes for which upregulation is amplified by the DHA treatment (n=138); “amp down” -genes for which their IDC-mediated downregulation is accelerated by the DHA treatment (n=191) (**Figure 3B**, middle row). Finally, there are two DDEPs in which the natural IDC-mediated trend was decelerated by the DHA treatment, including “less up” genes whose baseline upregulation is attenuated (n=140); and “less down” genes whose baseline downregulation is decelerated (n=187) (**Figure 3B**, bottom row).

Together, these results suggest that the apparent reversal of the IDC transcriptional cascade in trophozoites by the DHA treatments reflects distortions in the individual transcriptional profiles of trophozoite-specific genes through either attenuating, accelerating, or reversing their baseline IDC-driven transcriptional profiles. To test if these patterns corresponded to the baseline transcription patterns in the IDC, we tested for enrichment of genes that show peak expression at specific points in the IDC in each expression pattern (**Figure 3C**). Indeed, genes whose transcription peaks at the ring stage are highly overrepresented in the “flip up” group. Genes with peak expression levels in the trophozoite stages are also overrepresented in the “flip down” and “amp down” DDEP. Overall, this indicates that the DHA-induced reversal of the IDC cascade from trophozoite to a ring-like transcriptional profile is mainly driven by the suppression of late-stage transcripts and the reinduction of early-stage transcripts. Indeed, the reduction of trophozoite stage parasite after 6 hr exposure to 150nM DHA in the trophozoite treatment course was also confirmed by microscopy (**Figure S8 & S9**)

### Transcriptional coregulation of protein complex components, including the Kelch-interacting complex (KIC)

Transcriptional co-regulation of genes encoding interacting proteins, such as protein complex subunits, is typically highly coregulated through various species, including *S. cerevisiae*, *D. melanogaster*, *H. sapiens*, *C. elegans*, and *A. thalian*a^50–53^. There is also evidence that at least some protein-protein interacting pairs have correlated mRNA abundance profiles in *P. falciparum*^54^. Co-regulation analysis of dose-responsive transcripts, or those that share a DDEP, could provide greater granularity in the parasite’s functional response to DHA, as functional enrichment yielded only general terms and components of protein complexes (**Supplementary Dataset 3**). To test this, we inspected the trophozoite DDEP results to find documented protein complexes that were overrepresented in each pattern. Specifically, we noted four significantly overrepresented gene groups forming putative *P. falciparum* protein complexes. These included PfKelch13 interacting proteins (e.g., KIC complex) ^19^ in the “flip up” group, condensin components in the “less up”, and proteasome and respiratory complex V components in the “flip down” (**Figure 4A-D**). For most (if not all) subunits of these four protein complexes, the change in their corresponding mRNA abundance over 6 hours (Δexp/6h) appeared highly concordant at each dose.

**Figure 4.**
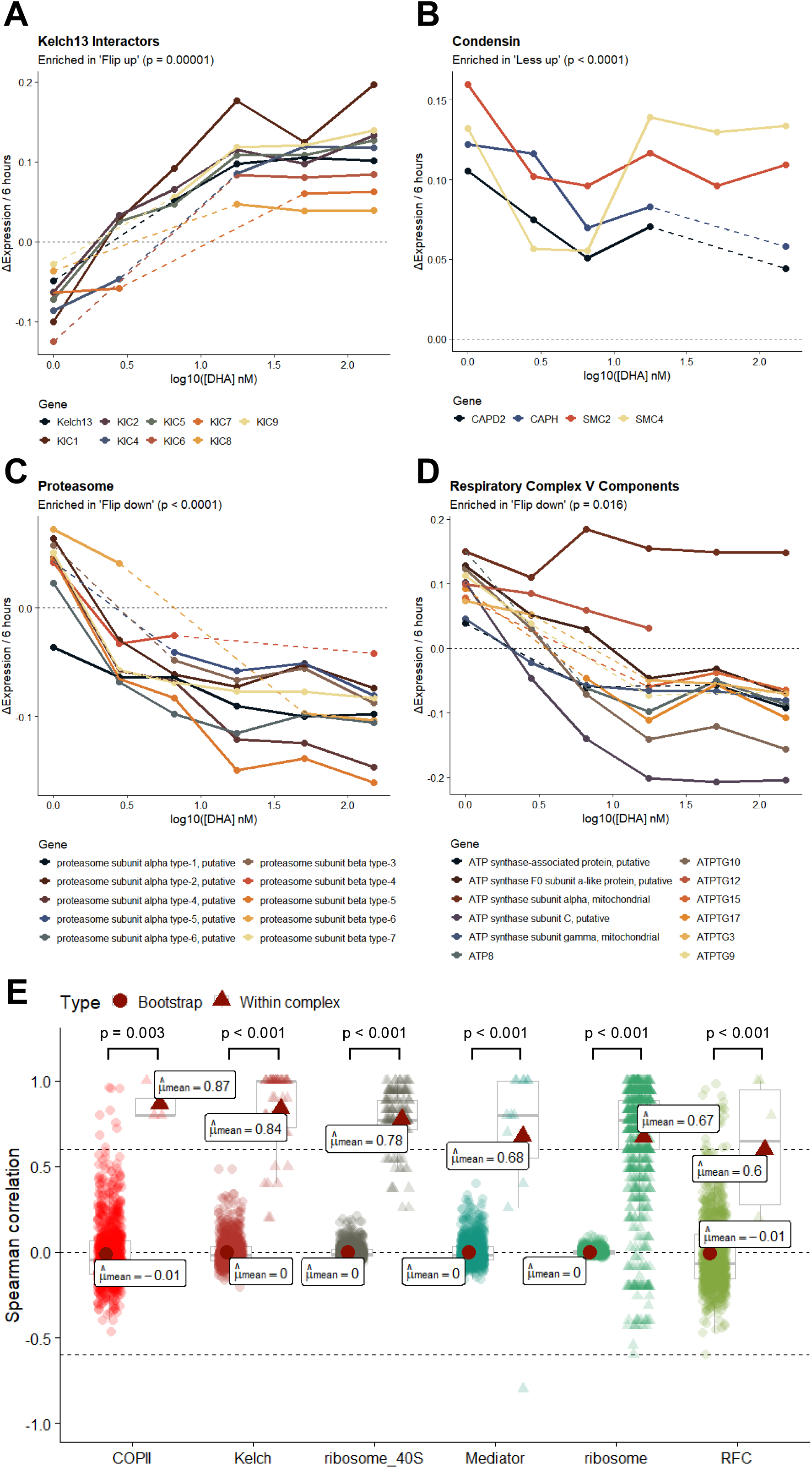
Coregulation of protein complex components. **(A-D)** Expression dynamics of protein complex components which were found to be enriched in trophozoite DDEPs: Kelch13 interactors **(A)**, condensin **(B)**, proteasome **(C)**, and respirator complex V **(D)**. Complex components that are enriched in the DDEs tend to show high similarity between the pattern of changes in expression over six hours. Points reflect the calculated change in expression between 0 and 6 hours of treatment at each dose in trophozoites. Dashed lines represent linearly interpolated values between points when the slope of an intermediate dose could not be calculated. P-values indicate adjusted hypergeometric p-values by Benjamini-Hochberg method. **(E)** Distribution of correlation values (y axis) of 6-hour change in expression across doses in protein complex components (x axis). Triangles show pairwise correlation of complex components, while circles show the null distribution generated by bootstrap testing. The global mean of each distribution is annotated with a large dark red circle (null distribution) or a dark red triangle (within complex). The difference in population means is tested by the Wilcoxon signed-rank test and significant p-values are indicated above the distribution plots. Dotted horizontal lines denote cutoff correlation values and null correlation at 0.

To quantify this further, we calculated the pairwise Spearman correlation of Δexp/6h of each gene within a curated list of protein complexes recently supported by a Thermal Protein Profiling (TPP)-based methodology, MAP-X^55^. For each complex with *n* genes, this calculation was compared pairwise using Spearman correlations, calculated 1000 times with *n* genes randomly selected from the initial pool of 1643 genes, to obtain a null distribution (**Materials and Methods**). Here, we observed a statistically significantly increased correlation of pairwise Δexp/6h for several protein complexes with a within-complex correlation value of 0.6. These included the COPII, Kelch interacting partners (Kelch), mediator, replication factor C (RFC), 40S ribosome, and 60S ribosome, which showed significant within-complex correlation values (**Figure 4E**). These represent significant shifts in mean within-complex correlation from the null distribution mean by the Wilcoxon signed-rank test. Most striking is the high correlation (mean ρ = 0.84) between the change in transcript level of PfKelch13 and seven of its interacting partners, KIC1 (PF3D7_0606000), KIC2 (PF3D7_1227700), KIC4 (PF3D7_1246300), KIC6 (PF3D7_0609700), KIC7 (PF3D7_0813000), KIC8 (PF3D7_1014900), and KIC9 (PF3D7_1442400). These have been found to share an endocytic compartment, and their coordinated transcriptional induction by DHA corroborates their significance to the parasite’s response to DHA. In addition, condensin, exosome, minichromosome maintenance (MCM), microtubules, proteasome regulation, respiratory complex III, and T-complex also exhibited a trend of increased correlation of their pairwise Δexp/6h correlations, albeit they did not pass the 0.6 correlation threshold (**Figure S10**). This observation suggests that for some protein complexes, transcriptional regulation of their corresponding subunits is highly orchestrated and responsive to external stresses (e.g., DHA treatment).

### Motif enrichment analysis

It is now well established that Cis-regulatory elements play an important role in transcriptional regulation of the *P. falciparum* transcriptional cascade, some of which are recognized by members of the AP-2 transcription factors ^56–59^. Up until now, however, no cis-regulatory elements have been associated with transcriptional responses of *P. falciparum* to external perturbations. The search for such regulatory elements is likely hindered by the complexity of the mRNA abundance profiles in *P. falciparum,* which are typically a composite of multiple events, combining specific transcriptional inductions/suppressions with the temporal progression of the IDC transcriptional cascade, as shown in this study. Here, we take advantage of the thorough transcriptional deconvolution conducted in this study and search for putative *cis*-regulatory elements driving these transcriptional responses. We utilize the gene grouping associated with individual DDEPs and with transcripts of protein complexes that share DDEPs. For that, we examined the presence of 8bp motifs within the 1000bp upstream of transcription start sites with the *de novo* motif discovery tool FIRE^60^. We found three motifs to be enriched in the “flip down” DDEP and one motif enriched in the “flip up” DDEP, two of which are located upstream of the genes and one within the 3’UTR (**Figure 5A**). Since the “flip down” DDEP corresponds to late-stage transcripts, these motifs may either signal activation of late-stage transcripts or suppress their transcription during early stages. Conversely, the motif enriched in “flip up” genes may either induce early-stage transcripts or suppress them as parasites progress to the trophozoite stage. As such, the native role of at least some of the identified motifs may be linked with the regulation of the *P. falciparum* IDC transcriptional cascade and only altered by external perturbations.

**Figure 5.**
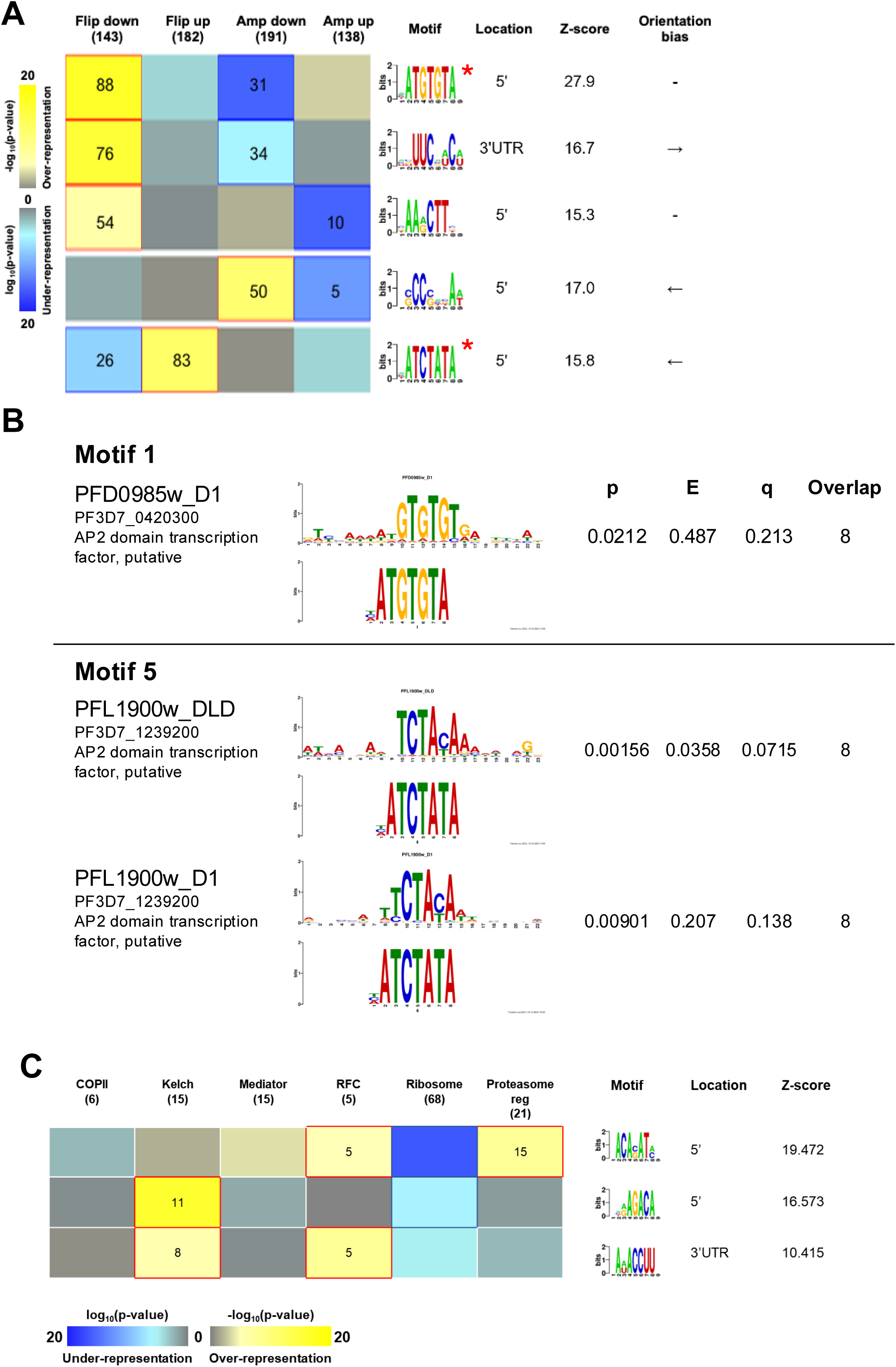
Motifs enriched in genes sharing DDEPs. (A-B) *De-novo* discovered motifs enriched in A) DDEPs with pattern reversal or pattern acceleration or B) protein complexes with high correlation in gene changes. Colors in the heatmaps correspond to p-value for enrichment of a motif in the 1000bp upstream of TSS of a gene in the group of genes input per column, with yellow for over-representation and blule for under-representation. The numbers in headers indicate the number of genes input, while the numbers in colored cells indicate the number of genes having the corresponding motif. Red outlines indicate statistically significant p-values. **(C)** Motif similarity analysis between two *de novo* motifs discovered in **(A)** and *in vitro* AP2 transcription factor binding sites discovered by Campbell *et al.* (2010).

To test this, we sought similarities between the identified motifs and known transcription factor binding sites. We compared the *de novo* motifs to the AP2 binding sites reported by Campbell, L.T. *et al* (2010)^56^ using the Tomtom Algorithm^61^. Indeed, Motif 1, which was over-represented in the “flip down” DDEP, had significant similarity to the binding site of PF3D7_0420300, a putative AP2 transcription factor (**Figure 5B**, top panel). Motif 5, which was enriched in the “flip up” DDEP, was similar to two binding sites of PF3D7_1239200, another putative AP2 transcription factor (**Figure 5B**, bottom panel). Due to the high correlation among DDEPs of protein complex components, we also tested for motif enrichment in differentially expressed genes within those complexes (**Figure 5C**). Two motifs were found enriched in Kelch-interacting genes, one of which was also enriched in replication factor C (RFC) components. Another motif was enriched in the proteasome regulatory complex and was also present in RFC genes.

## Discussion

By far, this study is not the first exploration of transcriptional responses of *P. falciparum*, the deadliest malaria parasite species, to artemisinin, the main component of the current frontline chemotherapy. The artemisinin-induced transcriptional dysregulation of numerous pathways that we observed in this study has indeed been reported earlier, suggesting the capacity of *P. falciparum* to respond to the cytotoxic effect of artemisinin. Artemisinin is believed to exert its effects through nonspecific or semispecific alkylation of numerous parasite proteins^62^, leading to proteotoxic^63^ and oxidative stress^64^ and destabilization of protein complexes, including the ribosomes^65^. Related to this concept, previous studies have reported transcriptional alterations in key metabolic and cellular pathways, including protein biosynthesis and ribosome biogenesis, protein turnover, cellular adhesion, lipid metabolism, and protein transport^21,29,34–39^. Collectively, these studies provided strong evidence for the transcriptomic responsiveness of each individual pathway to artemisinin. However, little was known about the context of these responses within the progression of the IDC, their dose-responsive character, and their potential epistatic relationships that determine artemisinin efficacy during the P. falciparum asexual blood stages (e.g., IDC), ^2,3^. Here, we showed that the majority of the transcriptional response occurs in the middle of the IDC, the trophozoite stage, including most, if not all, of the abovementioned artemisinin MOA-related pathways. The fringes of the IDC, rings and schizonts, on the other hand, possess much lower capacity of responsive in gene expression regulation, the earliest of which (rings) is particularly limited. This correlated with parasites insensitivity at later ring stage and schizont stage^6^. This result is consistent with the perceived key MOA component of artemisinin, which affects protein synthesis and homeostasis and whose factors are most transcriptionally active in trophozoites^46^. Indeed, the broad downregulation of these pathways (**Figure 2H**) that otherwise dominate the transcriptional profile of the trophozoites, is reminiscent of the reversal of the IDC transcriptional cascade back towards the ring stages. This observation yields a speculation that sensing the disruptions in protein homeostasis, the *P. falciparum* parasite has the ability to downregulate its protein synthesis, which coincides with the progression from ring to trophozoite stages. This reversal could also be linked to the induction of a previously documented dormant parasite stage, characterized by complete suppression of protein synthesis as an overall transcriptional program carrying hallmarks of quiescence and senescence^66^. In future studies, it will be interesting to investigate the transcriptional regulation guiding this process, be it induced by specific effects of artemisinin and/or other general stress response mechanisms shared with other external perturbations such as heat shock, which was recently related to the effect of artemisinin in a large-scale saturation mutagenesis screen^67^.

Transcriptional dysregulation of factors of protein synthesis and protein homeostasis/turnover is also strongly linked with the *P. falciparum* artemisinin resistance phenotype. First detected in Southeast Asia in 2008^8^, *P. falciparum* infections with slow-clearing parasitemia are now progressively emerging in Sub-Saharan Africa, signalling the loss of full effectiveness of ACT in the near future in the malaria endemic region^10^. Transcriptional reprogramming was observed already in the first analyses of the transcriptional profile of the parasites with this so-called partial artemisinin resistance phenotype (PARP), which were conducted with the *P. falciparum* cohort from the Tracking Artemisinin Resistance Collaboration (TRACI) project across the Greater Mekong Subregion (GMS) between 2011 and 2013^68^. Investigating the in vivo transcriptome of ∼1000 *P. falciparum* parasites in a blood sample showed a marked upregulation of processes associated with protein synthesis and turnover in the resistant parasites, concomitant with a deceleration of the IDC^69^. The latter was also confirmed *ex vivo*^49^. Indeed, transcriptomic analyses of a subsequent TRACII cohort of *P. falciparum* from uncomplicated malaria infections collected across the GMS between 2016 and 2018, revealed that the transcriptomic response of *P. falciparum in vivo* resembles the one *in vitro*, with marked downregulations of factors of not only protein synthesis but also hemoglobin digestion, energy metabolism, and membrane homeostasis^24^. This was, however, observed only in susceptible parasites. The resistant parasites analyzed in TRACII showed no transcriptional responses and instead appeared to exhibit a constitutively expressed profile (artemisinin resistance-associated transcriptional profile, ARTP) that resembled inducible changes in susceptible parasites. Moreover, the ability of *P. falciparum* to decelerate IDC progression following artemisinin exposure was demonstrated in the TRACII cohort of *P. falciparum* natural infections and subsequently in an additional *in vivo* cohort (TACT-CV) collected in the western and eastern provinces of Cambodia between 2018 and 2020^22^. The latter study was particularly significant, demonstrating that the susceptible parasites undergo a massive transcriptional downregulation of genes associated with protein synthesis as well as trophozoite-linked functionalities, which is, however, not the case for the PARP-carrying parasites^22^. Altogether, these results suggest that *P. falciparum* parasites with PARP are characterized by a constitutive transcriptional reprogramming of multiple factors/genes, some of which were initially involved in an inducible transcriptional response to the artemisinin parasiticidal activity. This was also confirmed by *in vitro* analysis of *P. falciparum* with non-synonymous SNPs in the *pfkelch13* gene, the main marker of artemisinin resistance^21^. There is emerging evidence that at least some of the reprogramming of these genes is mediated by genetic polymorphisms in their promoters or other DNA regulatory regions that evolved in the PARP parasites under drug pressure^23^. One of these factors is the *P. falciparum* cyclophilin 19B (PfCYP19B), whose transcriptional levels are controlled by sequence-length polymorphism and a dinucleotide (AT) short tandem repeat (STR) in its promoter region^70^. In future studies, it will be interesting to explore genetic polymorphisms of *P. falciparum* promoter regions in the context of artemisinin resistance (e.g. Genome Wide Association Studies, GWAS). Evidence from this study, linking individual genes and biological processes to artemisinin MOA, could guide these studies further. Focusing on sequence-length polymorphisms of STRs, which were recently shown to play a role in the evolution of *Plasmodium* parasite populations, is particularly attractive^71^.

To our knowledge, this study is the first to apply dose-responsive transcriptomics combined with time-course analysis to study *P. falciparum* response across multiple IDC stages. It is reasonable to suggest that, with this approach, one can explore transcriptional responses of malaria parasites to external perturbations in a higher-level information context, resolving the dilemma between a hard-wired transcriptional profile and phenotypic plasticity mediated by gene expression regulation in *Plasmodium* parasites, as debated earlier^30,31^. Here we show that under certain circumstances (e.g., IDC stage, time, and concentration exposures) *P. falciparum* is barely responsive to external perturbation, such as early ring stage after even the highest concentration of DHAat 6hr. Under presumably favorable conditions, such as the trophozoite stage at a range of DHA concentrations, can induce broad transcriptional changes that are both directly inducible at short intervals (1hr) and embedded into the regulation of the IDC transcriptomics cascade at longer exposure (4-6hr). Moreover, the BMD distribution of dose-dependent DEGs closely mirrors fluctuations in DHA susceptibility throughout the IDC, as well as its clinical and other pharmacodynamic properties. As such, it is feasible to suggest that similar comprehensive transcriptomic analyses can be applied to future phenotypic drug discovery, agnostic to molecular targets^72^. These could be particularly useful in assessing the MOA of experimental compounds, such as those obtained by massive phenotypic screening efforts, which have yielded thousands of hits, and the MMV Malaria Box compound library^73–75^. Results from these studies have the potential to bring transcriptome profiling back into the “toolbox” of system biology methods, as a scalable approach that produces comprehensive data on biological changes in response to drug treatment. Interpretation of transcriptomic profiling experiments such as this study greatly benefits from databases that allow comparison of profiles resulting from similar or contrasting perturbagens to construct hypotheses regarding their MOA^72^.

## Materials and Methods

### In vitro parasite culture and synchronization

A) P. falciparum strain 3D7 (MR4) was cultured according to standard methods^76^. Briefly, cultures were maintained in 2% hematocrit of human red blood cells (Innovative Research, USA) in malaria culture medium (MCM) consisting of RPMI 1640 (Gibco), 0.25% (w/v) Albumax II (Gibco), 2g/L sodium bicarbonate (Sigma), 0.1 mM hypoxanthine dissolved in sodium hydroxide (Merck), and 50 μg/L gentamycin (Gibco). Cultures were incubated at 37°C in a gas mixture of 3% oxygen, 5% carbon dioxide, and 92% nitrogen. Culture media was refreshed every 24-48 hours, and parasitemia assessed regularly by preparation of thin blood smears stained by 10% (v/v) Giemsa (Sigma). Parasite synchronicity was adjusted to a 6-hour window with 5% (w/v) D-Sorbitol (Sigma) and Percoll gradient centrifugation as previously described^77^.

### DHA treatment

Serial dilutions of DHA (WWARN) dissolved in DMSO (Sigma) were prepared with a top concentration of 150 nM and three-fold dilution in MCM, yielding final treatment concentrations of 150 nM, 50 nM, 18.7 nM, 5.6 nM, and 1.9 nM, with a 0 nM control. Parasite cultures were adjusted to between 3-5% parasitemia and treated at 2% hematocrit. For experiments in the ring stage, the experiment was conducted on rings between 8 hpi. The trophozoite-stage experiments were conducted at 16 hpi, and the schizont-stage experiments at 36 hpi. Parasite age assessment was conducted by microscopy of a Giemsa smear.

### RNA extraction

RNA was processed according to previously described methods^42^. For every volume of infected RBC to be harvested, 10 volumes of Trizol (Invitrogen) were added. The mixture was vortexed for 30 seconds and allowed to incubate at room temperature for 5 minutes and frozen at -80°C until RNA purification. Trizol mixtures were centrifuged for 1 minute at 15,000 RPM on a benchtop centrifuge at room temperature. The supernatant was removed and used as input for RNA extraction.

RNA purification was done with ZymoBIOMICS Magbead RNA kit (Zymo) either by hand or with an in-house liquid handler protocol (Hamilton). Magnetic bead purification was done on the Magnum FLX magnetic plate (Alpaqua). All steps up to elution followed manufacturer recommendations, where samples were eluted in 15 μL elution buffer (Zymo) instead of the recommended volume to preserve high RNA concentrations. DNAse treatment was not performed on any samples.

Purified total RNA was then quantified by the Quant-iT™ RNA High Sensitivity kit (Invitrogen) by automated liquid handler. RNA was then normalized with nuclease-free water. Samples with sufficient concentration were normalized by automation, while low-concentration samples were normalized manually. The target RNA concentration was typically set to 10 ng/μL or matched to the lowest sample concentration of that batch, with a minimum of 4 ng/μL. After normalization, the RNA Integrity Number (RIN) of 2-3 randomly selected samples per treatment condition was assessed by Agilent Bioanalyzer (Agilent).

### Library preparation

For preparation of non-stranded libraries, total RNA was reverse-transcribed by the SMART-Seq2 method as previously described^42^. In brief, to 5 μL of normalized mRNA, 4 μL of 1 μM Oligo-dT_20_VN and 1 mM dNTP mix was added and incubated at 72 °C for 3 minutes, then on ice for 5 minutes. Reverse transcription of each sample was done with 100U SuperScript II reverse transcriptase, 10 U RNaseOUT (Invitrogen), 1x SuperScript II first strand buffer (Invitrogen), 5 mM DTT (Invitrogen), 1 M betaine (Thermo Fisher), 6 mM MgCl_2_ (Sigma), 1 μM LNA-TSO (IDT DNA) topped up to 11 μL with nuclease-free water. The reaction was mixed and subsequently incubated at 42 °C for 90 minutes, followed by 10 cycles of 50 °C for 2 minutes and 42 °C for 2 minutes, and finally 70 °C for 15 minutes.

Second strand synthesis was done with 1x KAPA HiFi HotStart Ready Mix (Roche) and 3 μM ISPCR oligo (IDT DNA) topped up to 20 μL with nuclease-free water. This was mixed with 5 μL of reverse transcription product and incubated at 98 °C for 3 minutes, followed by 19 cycles of 98 °C for 20 seconds, 56 °C for 15 seconds, and 64 °C for 6 minutes, and finally held for 64 °C for 5 minutes and cooled at 4 °C. The amplified cDNA was purified using Ampure XP beads (Invitrogen) as previously described. This cDNA was normalized to 1 ng/μL and libraries were then prepared with the Illumina Nextera XT Library Prep Kit (Illumina) according to manufacturer protocol. All library prep was done at half-volume to conserve reagents.

The library size and quality of randomly selected samples were assessed by Bioanalyzer (Agilent). Libraries were then shipped to Novogene, where quality control by Bioanalyzer (Agilent) was confirmed for all samples. Each sample was then sequenced on the Novaseq 6000 at a depth of 3 GB per sample.

### RNA-Seq preprocessing

Raw reads were trimmed using Trim Galore and Cutadapt^78^ with the following command:

trim_galore -q 20 – phred33 -a <I7 adapter> -a2 <I5 adapter> --stringency 5 – trim-n -e 0.2 -o ./1_trim – length 35 –paired 0_raw_reads/<SAMPLE name>.fq.gz

Low-quality base pairs were trimmed with Phred score cutoff of 20 (1 in 100 chances of incorrect base call). This score was set low to maximize the number of reads to align, as only canonical mRNA transcripts are captured during sample and library preparation. A reference file with sample and i7 / i5 index information was referenced to build this command to remove adapters and indexes. The stringency score was set at 15, while the error rate was set to 0.2. At this stage, reads shorter than 75bp after adapter trimming are discarded. A second round of trimming was applied for removal of universal sequencing primers, after which reads shorter than 35 bp were discarded. Trimmed non-stranded reads were mapped to the *P. falciparum* 3D7 reference genome from PlasmoDB release 55 using HISAT2^79^ and SAMtools^80^. The maximum intron length was set to 3000 to account for the short *Plasmodium* introns and prevent erroneous mapping of reads spanning chromosome lengths. Only paired and uniquely mapped reads were used for assembly and quantification with Stringtie^81^ Transcripts per million (TPM) values were calculated from the remaining raw counts.

Before differential expression analysis, all genes in the non-stranded dataset were subjected to low-expression filtering. Filtering was conducted on each stage separately, as we anticipated that stage-specific expression patterns would impact transcript detection. After this step, only genes with at least 1 TPM in at least 40 / 61 samples per stage were used.

### Determination of differentially expressed genes

Selection of dose-responsive differentially expressed genes was done using the DRomics package^43^. The pipeline used is available on GitHub or by request. Each parasite stage (ring, trophozoite, schizont) and timepoint (1 hour, 4 hours, or 6 hours) in this experiment was analysed separately for determination of the dose-response. Raw counts of the genes that passed low expression filtering were used as input. Genes were normalized by variance-stabilizing transformation method and selected for differential expression by the quadratic trend test with a false discovery rate (FDR) cutoff of 0.05. Selected genes were tested for fit against one of five models: linear, exponential, hill, Gauss-probit, and log Gauss-probit and genes without good fit to a model were discarded. All formulae and parameter information as presented in the DRomics publication are available at https://github.com/jboentoro/3D-omics. The BMD of each dose-responsive gene was then calculated by the BMD.zSD method to produce the final list of dose-responsive genes. Values from the BMD.x-fold method were not used in analysis. This process was repeated for each stage and timepoint.

### BMD clustering and gene enrichment analysis

BMDs as calculated with the BMD.zSD method were clustered using Gaussian mixed model analysis with the R mixtools package^82^. The relevant code for this analysis is available on GitHub (https://github.com/jboentoro/3D-omics). In brief, each set (stage and treatment duration) was analyzed for a bimodal (n=2) mixed distribution. For sets where most DEGs have low BMD (trophozoites at 4 hours and 6 hours of treatment), a cutoff of 25 nM was used. Groups of DEGs were then tested for enrichment of GO biological process separately by each stage, treatment duration, and BMD group.

Genes were tested for enrichment against the GO biological process (BP) and metabolic function (MF) database, as well as a manually curated list of protein complex components compiled by Pazicky *et al.*^55^. The hypergeometric P-value was calculated and adjusted with the Benjamini-Hochberg method with a cutoff of *p* < 0.05. This was done in both R and the built-in PlasmoDB Gene Ontology enrichment tool.

### Calculation of IDC transcriptome correlation

Calculation of correlation of experimental samples to the reference IDC transcriptome is important to confirm parasites are in the correct stage at the time the experiment is conducted. Furthermore, it reveals deviations in the life cycle transcriptome progression that are associated with drug exposure. For each stage, the median TPM value from 3 replicates was calculated. The Spearman correlation to the reference transcriptome ^42^ was then calculated by resampling 1000 times. Thus, 95% confidence intervals were calculated for the correlation of each sample to each reference timepoint and were plotted.

### Dynamic differential expression pattern determination

Dynamic differential expression (DDE) patterns were designed to measure the additional dimensionality introduced by timepoint data. To calculate the DDE, each gene was first fed through the DRomics pipeline, using time as the predictor variable and expression values in raw counts as the response variable. This process was repeated for each dose of DHA and the untreated control. The result was a modelled curve for each gene that followed a pattern from 0 to 6 hours of the experiment. Formulae for the fitted curves were then used to back-calculate expression values for each dose to facilitate plotting of each dose against time on the same figure. The final step in DDE calculation is comparing the change in slope between treated and untreated samples. Only the expression values at 6 hours were used in this calculation. Classification was then done by comparing change in the highest available dose for the gene with the 0 DHA sample.

Usage of the DRomics pipeline allowed filtering of noisy genes and those with no time-dependent pattern of expression. The limitation was that not every dose for each gene could be modelled against time, resulting in gaps in the data. For genes where the reference 0-dose could not be calculated, data was filled with reference IDC transcriptome data. As the magnitude of change is not comparable with our experimental data, this was only used as reference data when the direction of expression after treatment is opposite from untreated and was not used for the other DDEPs.

### Complex correlation DDE testing

To test for within-complex correlation of DDEs, a manually curated list of protein complex components compiled by Pazicky *et al.* was used as a reference^55^. For each dose, the 6-hour slope of each differentially expressed gene was calculated as previously described for each gene listed as a complex component. Modelled genes were removed from analysis if there was no trend detected at either 0, 1.8, or 150 nM DHA. Genes where the slope could not be calculated in fewer than 4 doses were also removed. Finally, complexes were excluded if the slope could not be calculated for more than half of the genes annotated as belonging to that complex. For each pair of genes within the complex, the Spearman rho of the 6-hour slope at each dose was calculated. Only complexes with a mean Spearman rho of at least 0.6 were tested for statistical significance.

As a negative control, bootstrap testing was performed. For every *n* genes in each tested complex, the same number of genes were randomly selected from all available genes. Pairwise Spearman rho values were calculated for these random genes and averaged. This process was then repeated 1000 times. To calculate statistical significance, the Wilcoxon signed-rank test was used to test differences in sample means between the within-complex and bootstrap groups.

### Drug susceptibility assays

Drug susceptibility of *P. falciparum* strains used in this study was assessed against dihydroartemisinin (WWARN) dissolved in DMSO (Sigma-Aldrich) using flow cytometry. Drug assays were set up to 2% hematocrit in 96-well plates with two-fold dilution of drug concentration and a control with no drug in two technical replicates. Starting parasitemia was set to 0.3-0.5% and plates were incubated for 60 hours under standard culture conditions, or until a full cycle had passed and the untreated control had reached 12-24 hours post-invasion. Spent media was then removed and cells were stained with SYBR Green (Invitrogen) and read on an Accuri C6 plus flow cytometer (BD). Flow cytometry results were gated and processed in C6 software (BD).

## SUPPLEMENTARY FIGURES

**Supplementary Figure S1.**
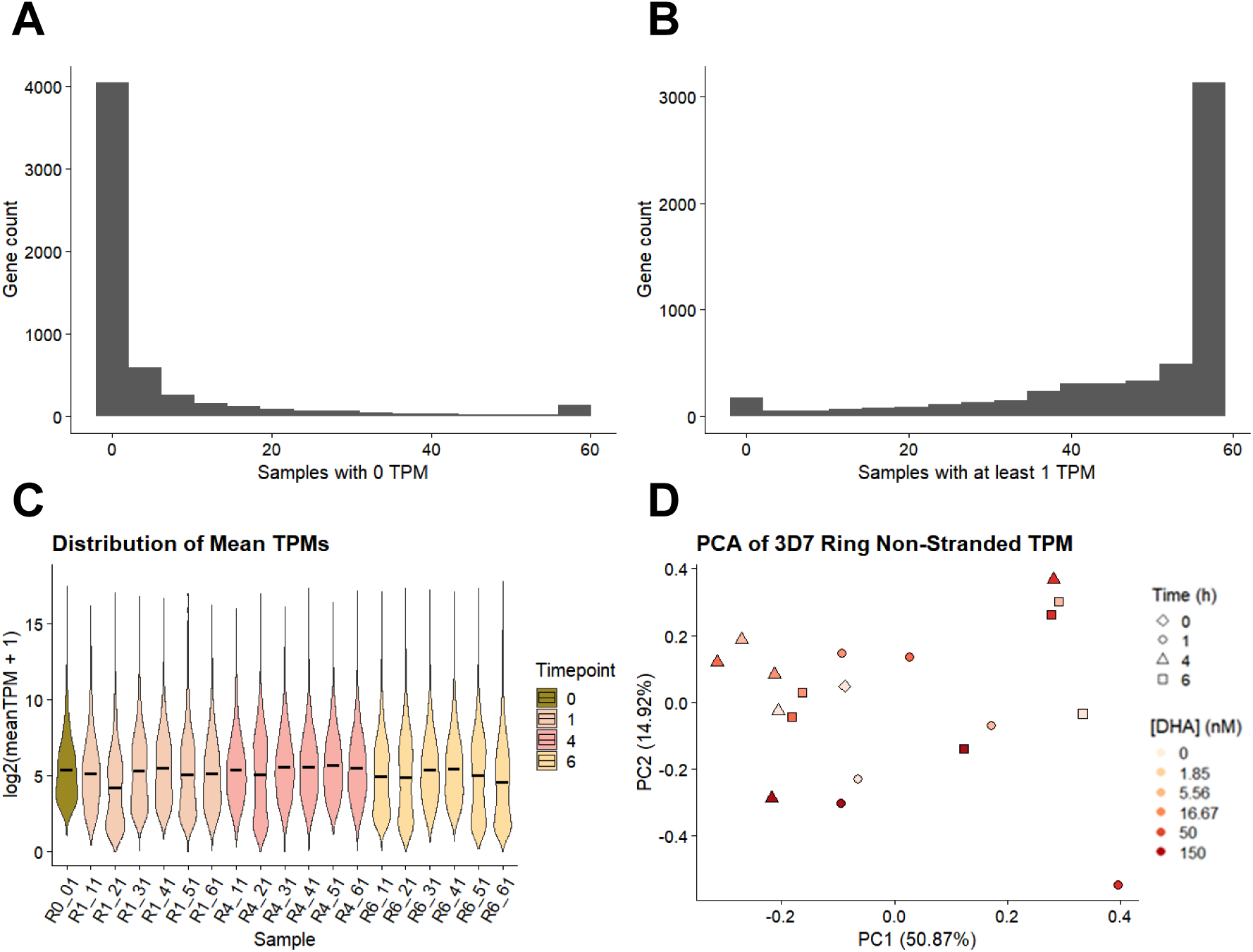
Expression filtering statistics in ring-stage samples. A) Histogram of the number of genes with 0 transcripts per million (TPM) (y-axis) in *n* number of samples (x-axis). B) Histogram of the number of genes (y-axis) with at least 1 TPM in *n* number of samples (x-axis). C) Violin plot showing distribution of log2(TPM +1) in each ring-stage sample. Colors denote treatment duration. Black crossbars indicate the mean of each group. D) PCA of ring-stage samples/ Shape denotes time of RNA harvest in hours after the start of treatment, while color indicates the dose of DHA.

**Supplementary Figure S2.**
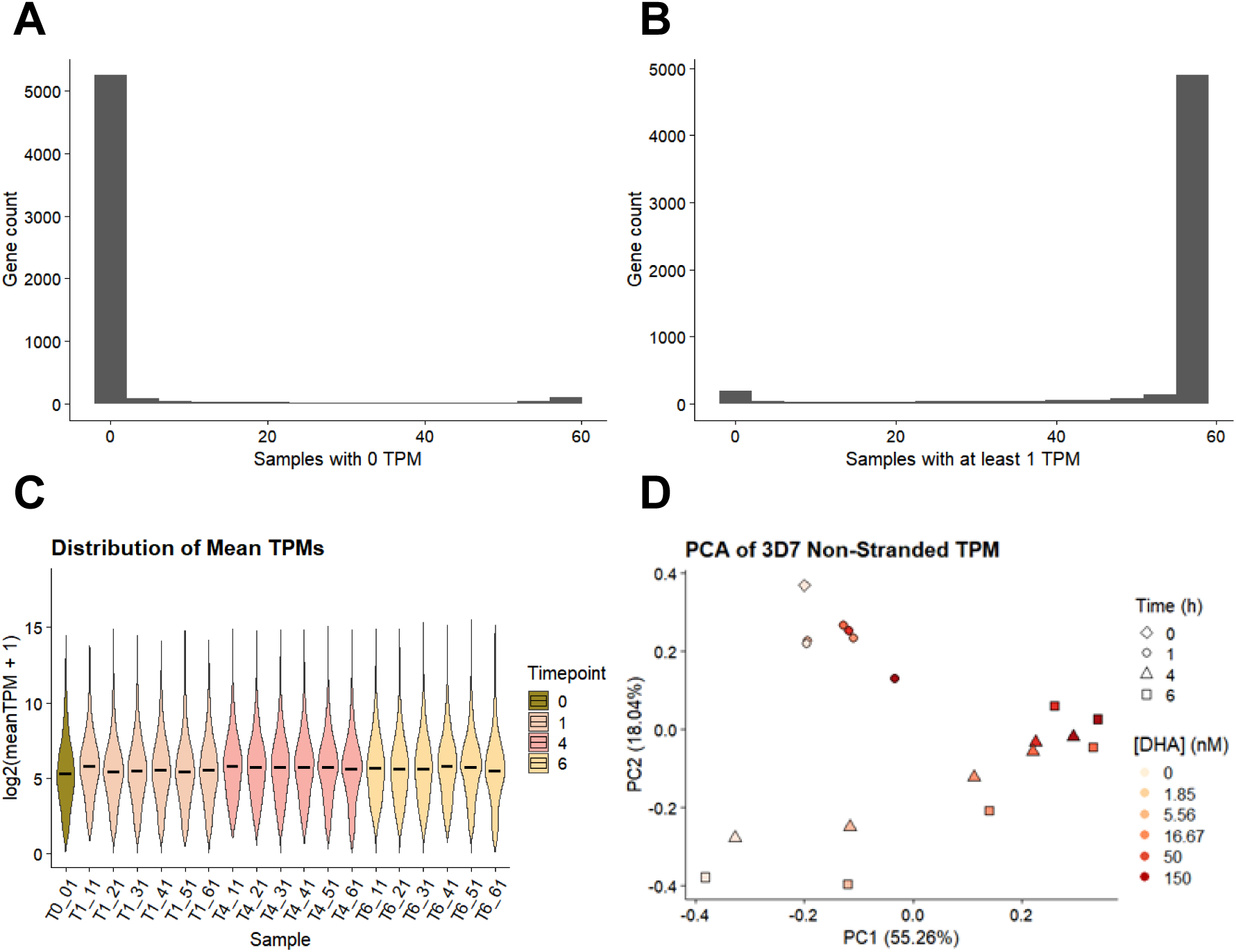
Expression filtering statistics in trophozoite-stage samples. A) Histogram of the number of genes with 0 transcripts per million (TPM) (y-axis) in *n* number of samples (x-axis). B) Histogram of the number of genes (y-axis) with at least 1 TPM in *n* number of samples (x-axis). C) Violin plot showing distribution of log2(TPM +1) in each ring-stage sample. Colors denote treatment duration. Black crossbars indicate the mean of each group. D) PCA of ring-stage samples/ Shape denotes time of RNA harvest in hours after the start of treatment, while color indicates the dose of DHA.

**Supplementary Figure S3.**
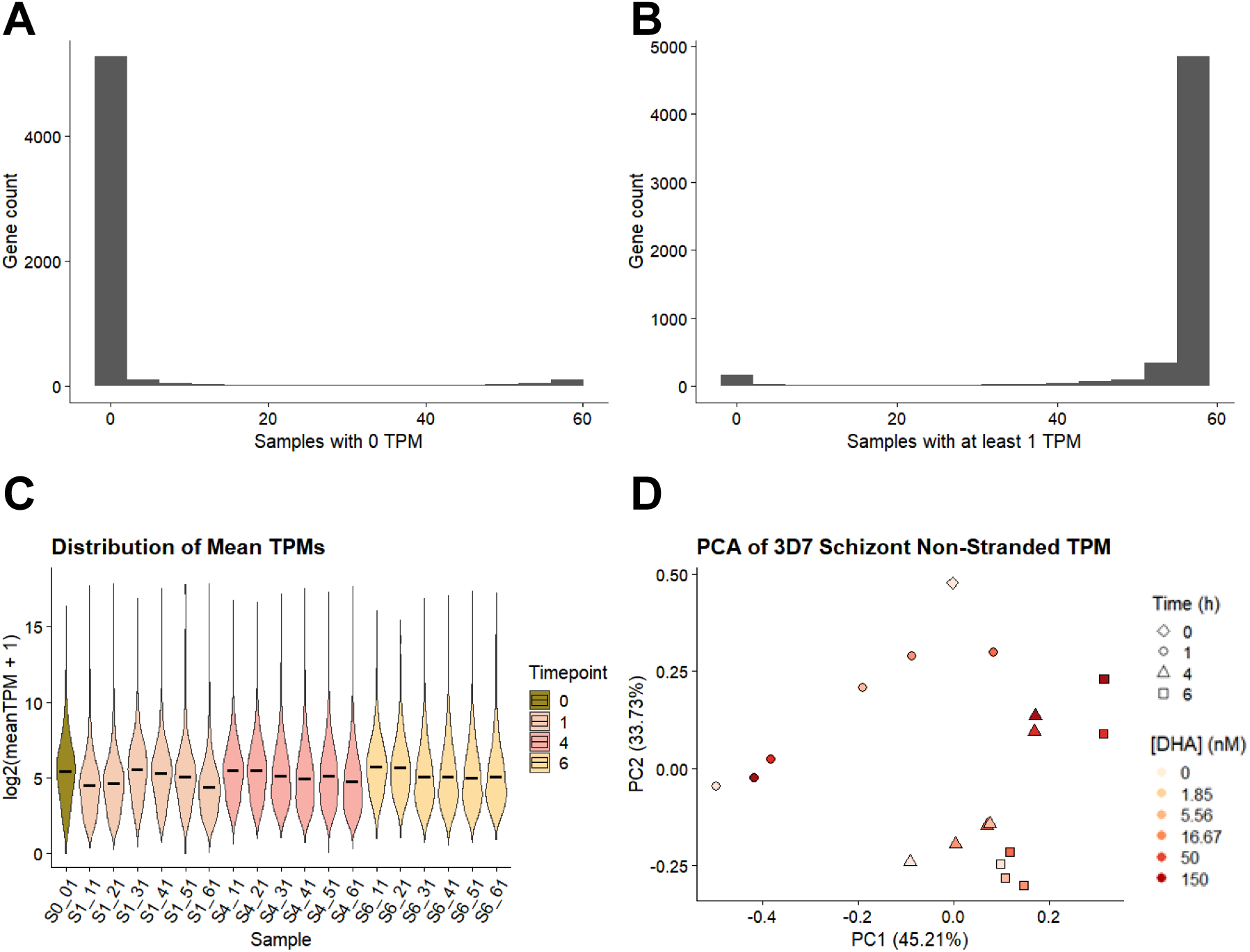
Expression filtering statistics in schizont-stage samples. A) Histogram of the number of genes with 0 transcripts per million (TPM) (y-axis) in *n* number of samples (x-axis). B) Histogram of the number of genes (y-axis) with at least 1 TPM in *n* number of samples (x-axis). C) Violin plot showing distribution of log2(TPM +1) in each ring-stage sample. Colors denote treatment duration. Black crossbars indicate the mean of each group. D) PCA of ring-stage samples/ Shape denotes time of RNA harvest in hours after the start of treatment, while color indicates the dose of DHA.

**Supplementary Figure S4.**
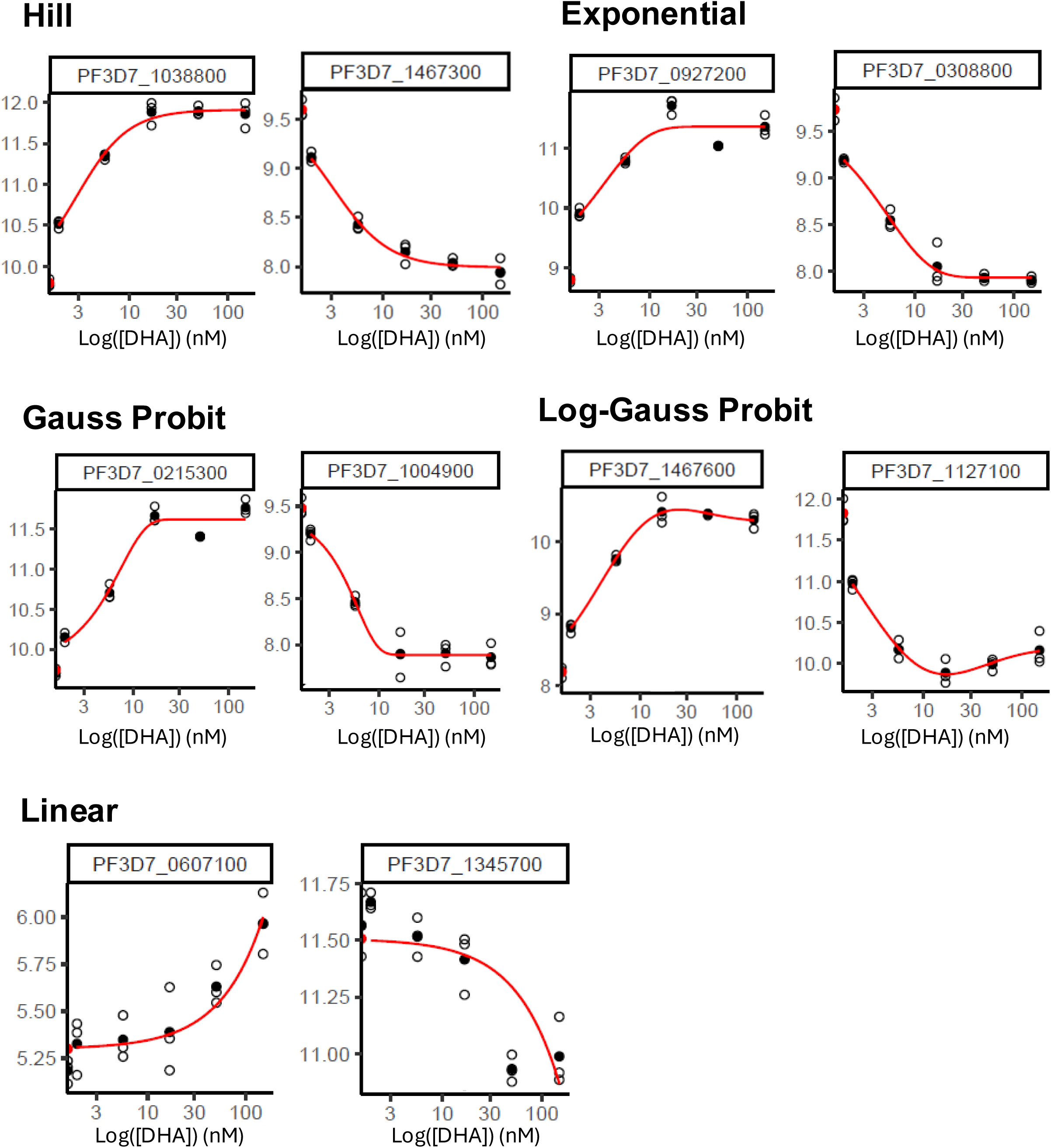
Examples of curves fitted to each model to determine dose-responsiveness. Output from DRomics R package.

**Supplementary Figure S5.**
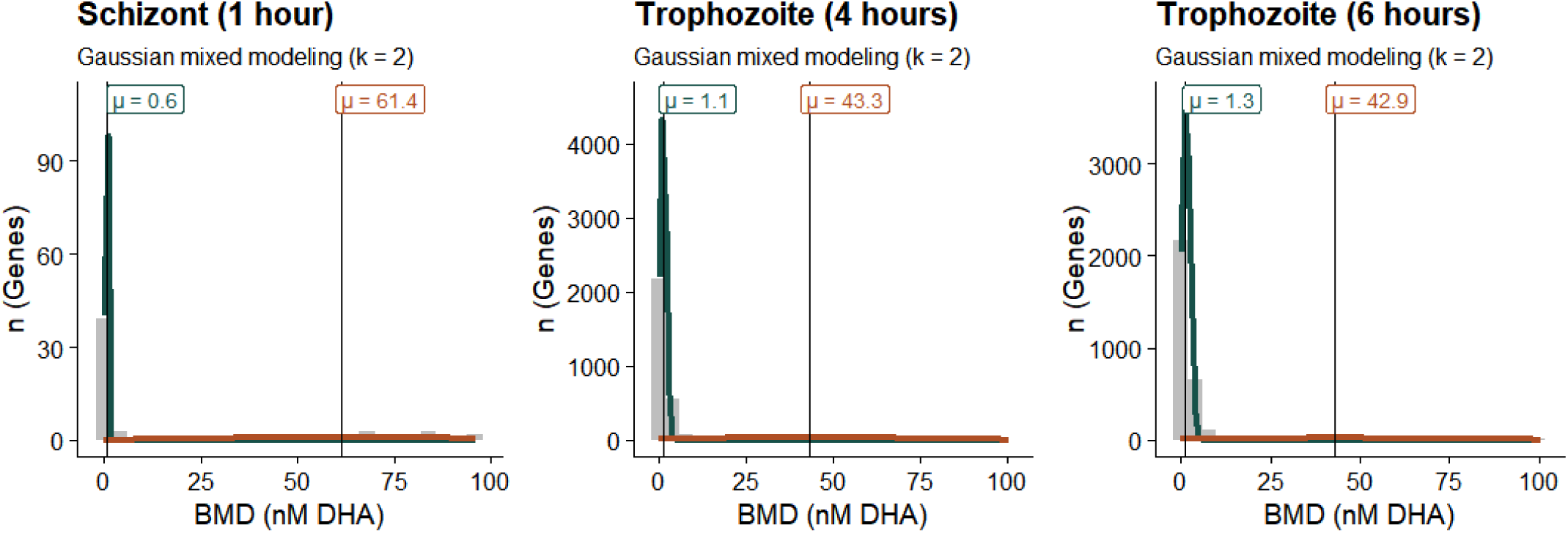
Gaussian mixed models of BMDs of DEGs. Description: Histograms of BMD values from schizonts treated for 1 hour and trophozoites treated for 4 and 6 hrs. The model in trophozoites failed to converged to a binomial model due to a much larger population with a BMDlow profile as compared to the BMDhigh profile. Solid lines represent the corresponding gaussian density distribution scaled to the gene counts. The line color corresponds to either BMDlow (green) or BMDhigh (brown). The mean of each distribution is annotated by a vertical line and text.

**Supplementary Figure S6.**
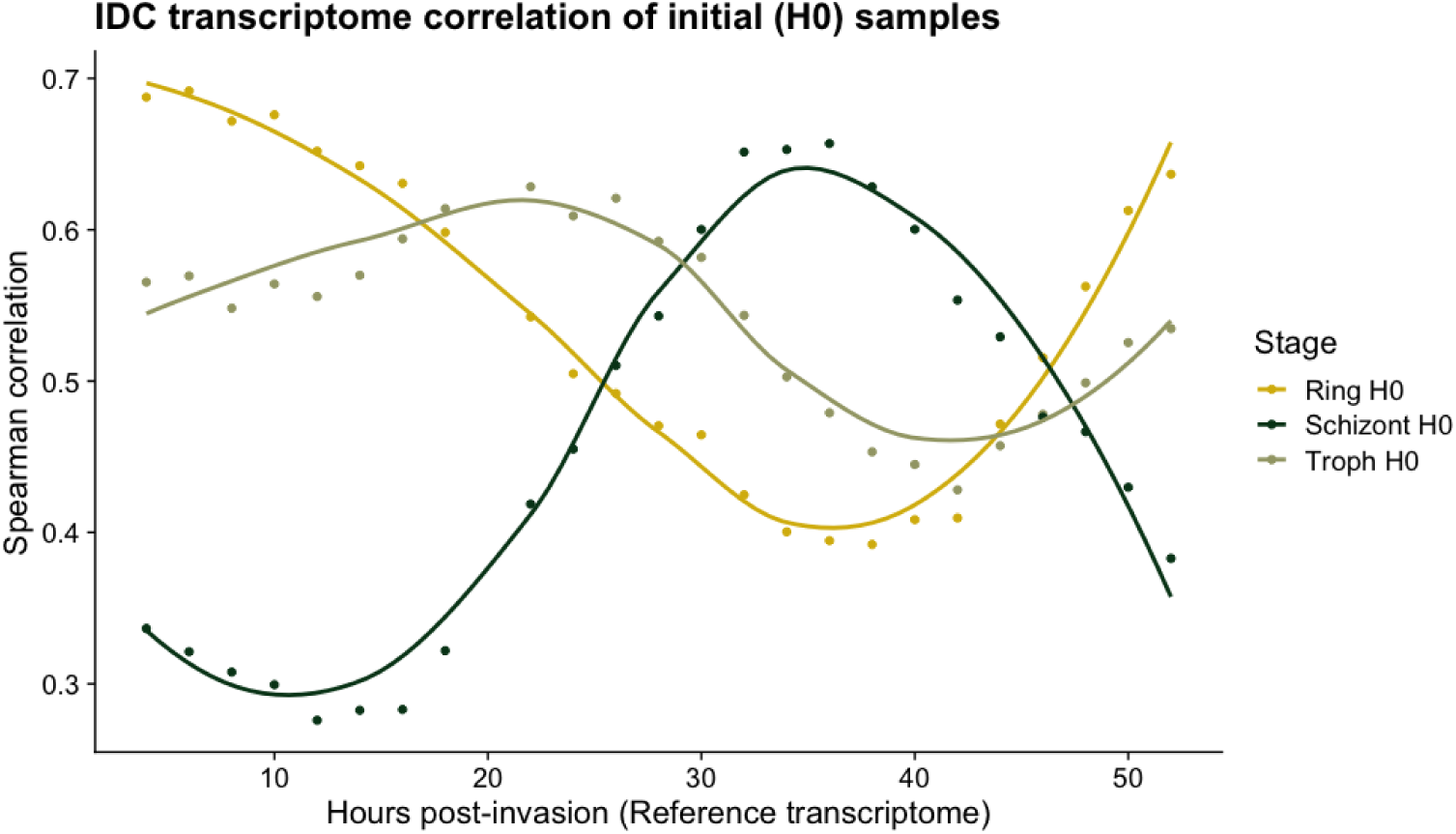
IDC transcriptome correlation of samples at the start of the experiment per stage

**Supplementary Figure S7.**
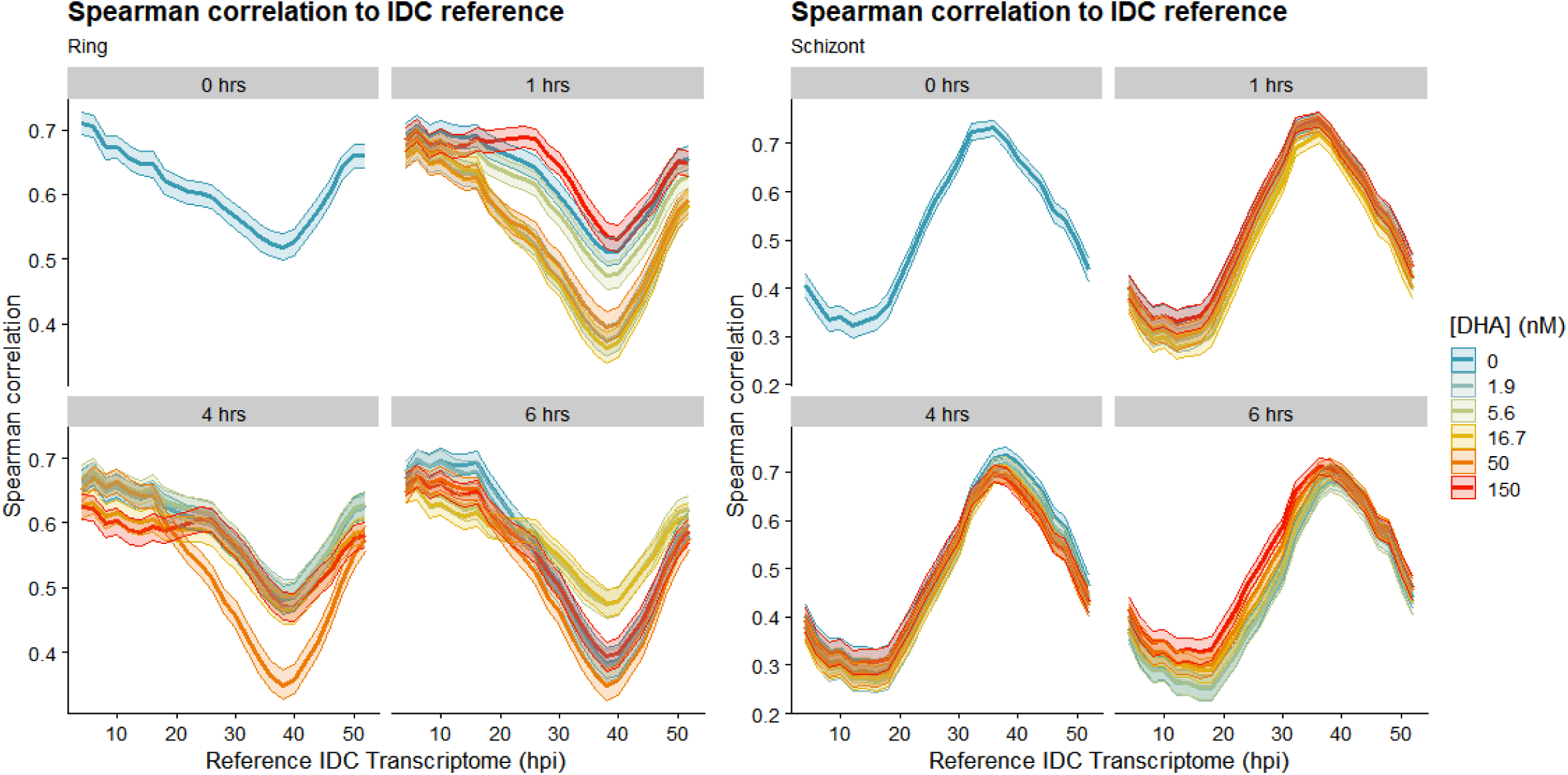
Spearman rank correlation heatmap of trophozoites during the experiment (grouped by treatment duration) with each sample in the reference IDC transcriptome generated by Kucharski et al. (2020). Row labels indicate the hours post-invasion (HPI) in the reference transcriptome dataset. Color scale indicates Spearman rank correlation. Green boxes delineate the range with maximum correlation to the untreated samples to indicate normal life cycle progression.

**Supplementary Figure S8.**
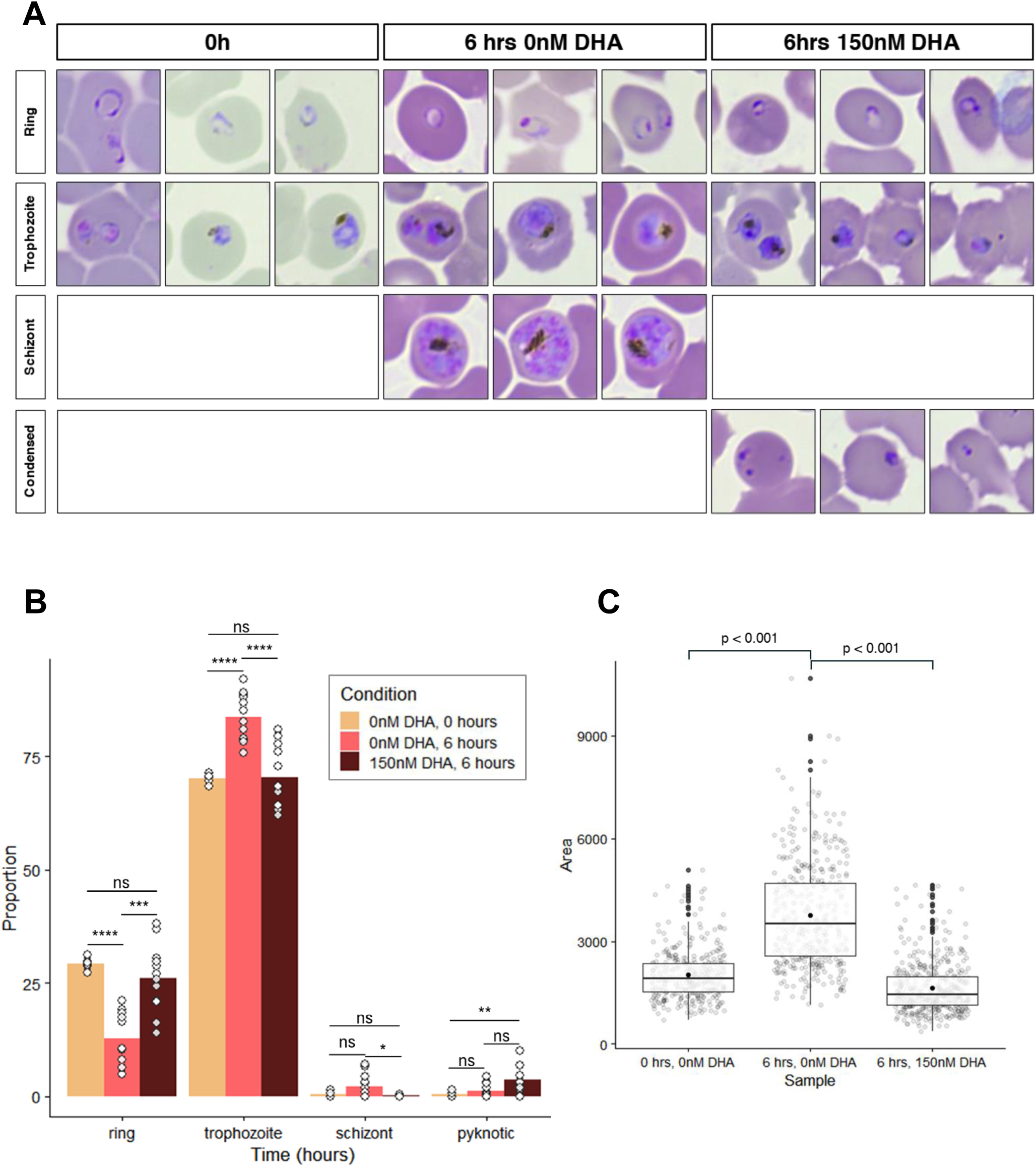
A) Representative parasite morphology at 0 and 6 hours with 150nM DHA (right column) or with 0nM DHA (middle column). These morphologies guided classification for calculation of proportion of parasite morphologies at each condition. B) Proportion of parasite morphology at each condition (combination of time and dose), counted from 3 biological replicates and 2 technical replicates by 2 independent microscopists. Bar height corresponds to the means of the morphology and condition tested, while points are the individual repetitions. Population means were compared by Brown-Forsythe and Welch ANOVA multiple comparisons, and statistically significant comparisons are marked with asterisks (** p<0.001, *** p<0.0001). C) Comparison of trophozoite area at each condition (n = 300) from microscopy images as measured by ImageJ. Population means were compared with a nested one-way ANOVA in GraphPad Prism 8.

**Supplementary Figure S9.**
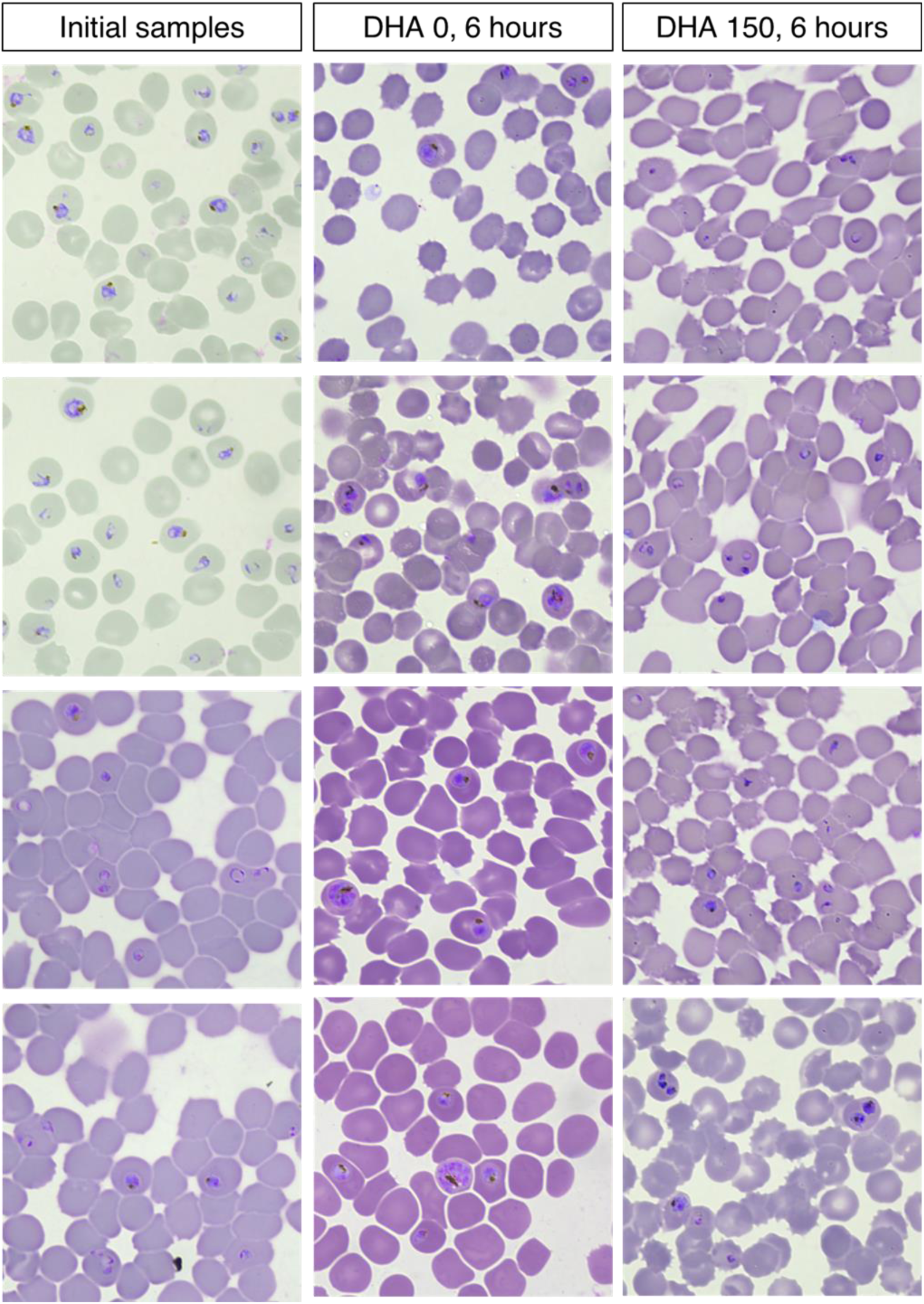
Representative microscopy fields of trophozoites a) before treatment (first column), b) at six hours without DHA (second column), and c) at six hours wih 150nM DHA. Smears were stained with 10% Giemsa or Field’s stain.

**Supplementary Figure S10.**
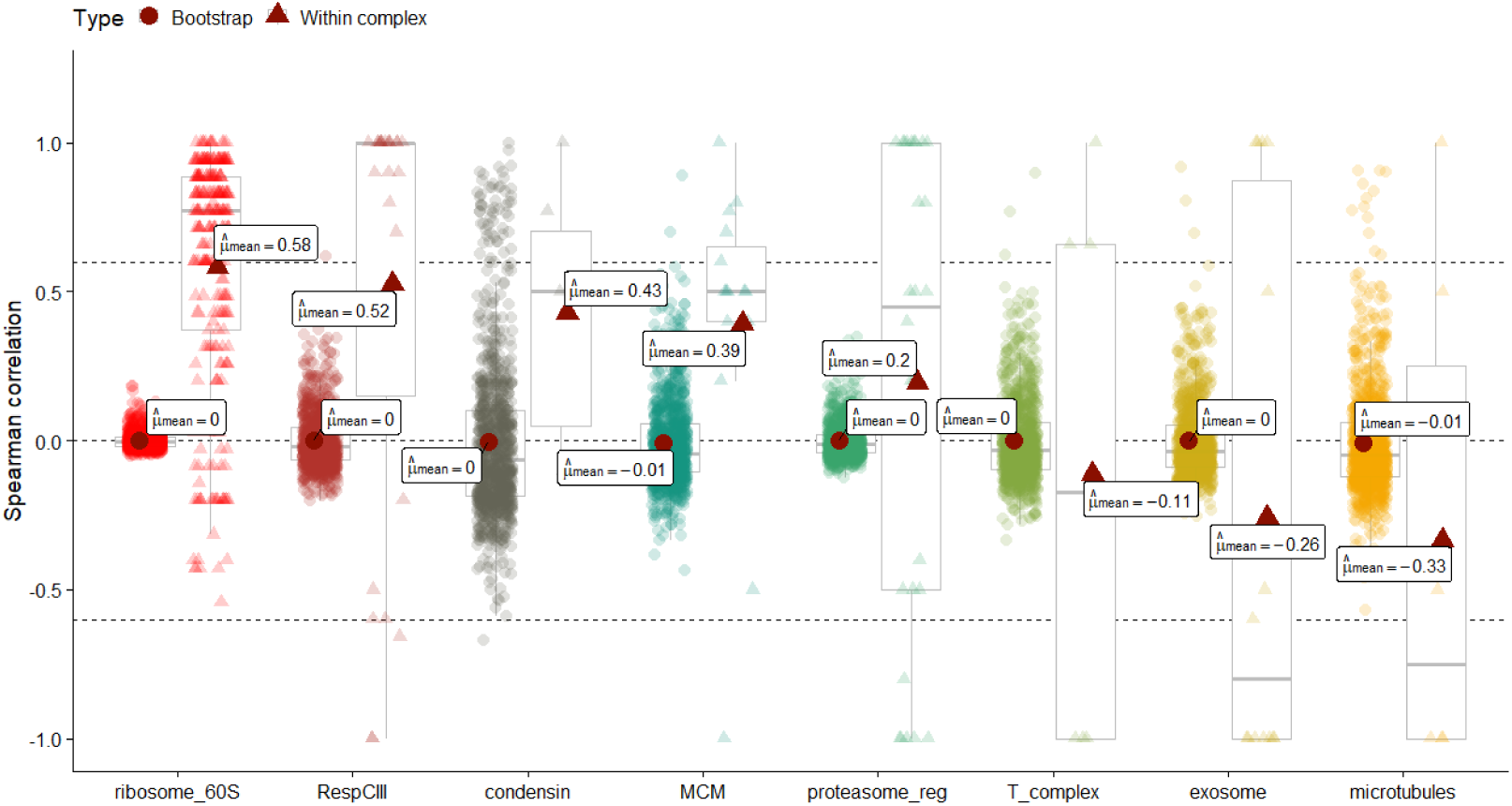
Protein complexes with non-significant dynamic differential pattern expression correlation in trophozoites (Mean rho < 0.6 and/or not significantly higher than the bootstrap mean).

## References

1. World Health Organization. World malaria report 2025: addressing the threat of antimalarial resistance. (Geneva, 2025). Licence: CC BY-NC-SA 3.0 IGO.

2. Aweeka, F.T. & German, P.I. Clinical pharmacology of artemisinin-based combination therapies. Clinical Pharmacokinetics 47, 91–102 (2008).

3. Cui, L. & Su, X.Z. Discovery, mechanisms of action and combination therapy of artemisinin. Expert review of anti-infective therapy 7, 999–999 (2009).

4. Antoine, T. et al. Rapid kill of malaria parasites by artemisinin and semi-synthetic endoperoxides involves ROS-dependent depolarization of the membrane potential. Journal of Antimicrobial Chemotherapy 69, 1005–1016 (2014).

5. Dogovski, C. et al. Targeting the Cell Stress Response of Plasmodium falciparum to Overcome Artemisinin Resistance. PLOS Biology 13, e1002132 (2015).

6. Klonis, N. et al. Altered temporal response of malaria parasites determines differential sensitivity to artemisinin. Proceedings of the National Academy of Sciences of the United States of America 110, 5157–5162 (2013).

7. Murithi, J.M. et al. Combining Stage Specificity and Metabolomic Profiling to Advance Antimalarial Drug Discovery. Cell chemical biology 27, 158–171.e3 (2020).

8. Dondorp, A.M. et al. Artemisinin Resistance in Plasmodium falciparum Malaria. New England Journal of Medicine 361, 455–467 (2009).

9. Balikagala, B. et al. Evidence of Artemisinin-Resistant Malaria in Africa. New England Journal of Medicine 385, 1163–1171 (2021).

10. Rosenthal, P.J., Asua, V. & Conrad, M.D. Emergence, transmission dynamics and mechanisms of artemisinin partial resistance in malaria parasites in Africa. Nature Reviews Microbiology *2024* 22:*6* 22, 373-384 (2024).

11. White, N.J. The parasite clearance curve. Malar J 10, 278 (2011).

12. Benoit, W., Didier, M., Chanaki, A. & Rick, M.F. Ring-stage Survival Assays (RSA) to evaluate the in-vitro and ex-vivo susceptibility of Plasmodium falciparum to artemisinins. 1–21 (Institute Pasteur du Cambodge _ National Institutes of Health Procedure RSAv1, 2014).

13. White, N.J. Emergence of Artemisinin-Resistant Plasmodium falciparum in East Africa. N Engl J Med 385, 1231–1232 (2021).

14. Dhorda, M. et al. Artemisinin-resistant malaria in Africa demands urgent action. Science 385, 252–254 (2024).

15. van der Pluijm, R.W. et al. Triple artemisinin-based combination therapies versus artemisinin-based combination therapies for uncomplicated Plasmodium falciparum malaria: a multicentre, open-label, randomised clinical trial. The Lancet 395, 1345–1360 (2020).

16. Ariey, F. et al. A molecular marker of artemisinin-resistant Plasmodium falciparum malaria. Nature *2013* 505:7481 505, 50-55 (2013).

17. Miotto, O. et al. Genetic architecture of artemisinin-resistant Plasmodium falciparum. Nature Genetics *2015* 47:*3* 47, 226-234 (2015).

18. Straimer, J. et al. K13-propeller mutations confer artemisinin resistance in Plasmodium falciparum clinical isolates. Science 347, 428–431 (2015).

19. Birnbaum, J. et al. A Kelch13-defined endocytosis pathway mediates artemisinin resistance in malaria parasites. Science 367, 51–59 (2020).

20. Mbengue, A. et al. A molecular mechanism of artemisinin resistance in Plasmodium falciparum malaria. Nature *2015* 520:7549 520, 683-687 (2015).

21. Mok, S. et al. Artemisinin-resistant K13 mutations rewire Plasmodium falciparum’s intra-erythrocytic metabolic program to enhance survival. Nature Communications *2021* 12:*1* 12, 1-15 (2021).

22. Nayak, S. et al. Population genomics and transcriptomics of Plasmodium falciparum in Cambodia and Vietnam uncover key components of the artemisinin resistance genetic background. Nature Communications *2024* 15:*1* 15, 1-17 (2024).

23. Zhu, L. et al. The origins of malaria artemisinin resistance defined by a genetic and transcriptomic background. Nature Communications *2018* 9:*1* 9, 1-13 (2018).

24. Zhu, L. et al. Artemisinin resistance in the malaria parasite, Plasmodium falciparum, originates from its initial transcriptional response. Communications Biology 5, 274–274 (2022).

25. Demas, A.R. et al. Mutations in plasmodium falciparum actin-binding protein coronin confer reduced artemisinin susceptibility. Proceedings of the National Academy of Sciences of the United States of America 115, 12799–12804 (2018).

26. Siddiqui, F.A. et al. Plasmodium falciparum Falcipain-2a Polymorphisms in Southeast Asia and Their Association With Artemisinin Resistance. The Journal of Infectious Diseases 218, 434–434 (2018).

27. Henrici, R.C., van Schalkwyk, D.A. & Sutherland, C.J. Modification of pfap2μ and pfubp1 Markedly Reduces Ring-Stage Susceptibility of Plasmodium falciparum to Artemisinin In Vitro. Antimicrobial agents and chemotherapy 64(2019).

28. Henriques, G. et al. The Mu Subunit of Plasmodium falciparum Clathrin-Associated Adaptor Protein 2 Modulates In Vitro Parasite Response to Artemisinin and Quinine. Antimicrobial Agents and Chemotherapy 59, 2540–2540 (2015).

29. Rocamora, F. et al. Oxidative stress and protein damage responses mediate artemisinin resistance in malaria parasites. PLoS pathogens 14(2018).

30. Hu, G. et al. Transcriptional profiling of growth perturbations of the human malaria parasite Plasmodium falciparum. Nature Biotechnology *2009* 28:*1* 28, 91-98 (2010).

31. Ganesan, K. et al. A Genetically Hard-Wired Metabolic Transcriptome in Plasmodium falciparum Fails to Mount Protective Responses to Lethal Antifolates. PLoS Pathogens 4, e1000214–e1000214 (2008).

32. Siwo, G.H. et al. An integrative analysis of small molecule transcriptional responses in the human malaria parasite Plasmodium falciparum. BMC Genomics 16, 1–17 (2015).

33. Otto, T.D. et al. New insights into the blood-stage transcriptome of Plasmodium falciparum using RNA-Seq. Molecular Microbiology 76, 12–12 (2010).

34. Asahi, H. et al. Profiling molecular factors associated with pyknosis and developmental arrest induced by an opioid receptor antagonist and dihydroartemisinin in Plasmodium falciparum. PLOS ONE 12, e0184874–e0184874 (2017).

35. Chen, N. et al. Fatty acid synthesis and pyruvate metabolism pathways remain active in dihydroartemisinin-induced dormant ring stages of plasmodium falciparum. Antimicrobial Agents and Chemotherapy 58, 4773–4781 (2014).

36. Oduor, C.I. et al. Single cell transcriptional changes across the blood stages of artemisinin resistant K13580Y Plasmodium falciparum upon dihydroartemisinin exposure. bioRxiv, 2023.12.06.570387-2023.12.06.570387 (2024).

37. Pires, C.V. et al. Chemogenomic Profiling of a Plasmodium falciparum Transposon Mutant Library Reveals Shared Effects of Dihydroartemisinin and Bortezomib on Lipid Metabolism and Exported Proteins. Microbiology Spectrum 11(2023).

38. Shaw, P.J. et al. Plasmodium parasites mount an arrest response to dihydroartemisinin, as revealed by whole transcriptome shotgun sequencing (RNA-seq) and microarray study. BMC Genomics 16, 1–14 (2015).

39. Tandoh, K.Z., Hagan, O.C., Wilson, M.D., Quashie, N.B. & Duah-Quashie, N.O. Transcriptome-module phenotype association study implicates extracellular vesicles biogenesis in Plasmodium falciparum artemisinin resistance. Frontiers in Cellular and Infection Microbiology 12, 886728–886728 (2022).

40. Barlow, S. et al. Guidance of the Scientific Committee on Use of the benchmark dose approach in risk assessment. EFSA Journal 7, 1150–1150 (2009).

41. Davis, J.A., Gift, J.S. & Zhao, Q.J. Introduction to benchmark dose methods and U.S. EPA’s benchmark dose software (BMDS) version 2.1.1. Toxicology and Applied Pharmacology 254, 181–191 (2011).

42. Kucharski, M. et al. A comprehensive RNA handling and transcriptomics guide for high-throughput processing of Plasmodium blood-stage samples. Malaria Journal 19, 363–363 (2020).

43. Larras, F. et al. DRomics: A Turnkey Tool to Support the Use of the Dose-Response Framework for Omics Data in Ecological Risk Assessment. Environmental Science and Technology 52, 14461–14468 (2018).

44. Crump, K.S. Calculation of Benchmark Doses from Continuous Data. Risk Analysis 15, 79–89 (1995).

45. Zhang, M. et al. Uncovering the essential genome of the human malaria parasite Plasmodium falciparum by saturation mutagenesis. *Science (New York*, N.Y.) 360, eaap7847-eaap7847 (2018).

46. Bozdech, Z. et al. The Transcriptome of the Intraerythrocytic Developmental Cycle of Plasmodium falciparum. PLOS Biology 1, e5 (2003).

47. Le Roch, K.G. et al. Discovery of gene function by expression profiling of the malaria parasite life cycle. Science 301, 1503–1508 (2003).

48. Zhu, L. et al. New insights into the Plasmodium vivax transcriptome using RNA-Seq. Scientific Reports *2016* 6:*1* 6, 1-13 (2016).

49. Mok, S. et al. Artemisinin resistance in Plasmodium falciparum is associated with an altered temporal pattern of transcription. BMC Genomics 12, 391 (2011).

50. Lee, J.W., Zemojtel, T. & Shakhnovich, E. Systems-level evidence of transcriptional co-regulation of yeast protein complexes. Journal of Computational Biology 16, 331–339 (2009).

51. Siwiak, M. & Zielenkiewicz, P. Co-regulation of translation in protein complexes. Biology Direct 10, 1–13 (2015).

52. Tan, K., Shlomi, T., Feizi, H., Ideker, T. & Sharan, R. Transcriptional regulation of protein complexes within and across species. Proceedings of the National Academy of Sciences of the United States of America 104, 1283–1288 (2007).

53. Webb, E.C. & Westhead, D.R. The transcriptional regulation of protein complexes; a cross-species perspective. Genomics 94, 369–376 (2009).

54. LaCount, D.J. et al. A protein interaction network of the malaria parasite Plasmodium falciparum. Nature *2005* 438:7064 438, 103-107 (2005).

55. Pazicky, S. et al. MAP-X reveals distinct protein complex dynamics across Plasmodium falciparum blood stages. Nat Microbiol 10, 3229–3244 (2025).

56. Campbell, T.L., de Silva, E.K., Olszewski, K.L., Elemento, O. & Llinás, M. Identification and genome-wide prediction of DNA binding specificities for the ApiAP2 family of regulators from the malaria parasite. PLoS pathogens 6(2010).

57. Quansah, E. et al. ApiAP2 Gene-Network Regulates Gametocytogenesis in Plasmodium Parasites. Cellular Microbiology 2022, 5796578–5796578 (2022).

58. Read, D.F., Cook, K., Lu, Y.Y., Le Roch, K.G. & Noble, W.S. Predicting gene expression in the human malaria parasite Plasmodium falciparum using histone modification, nucleosome positioning, and 3D localization features. PLOS Computational Biology 15, e1007329–e1007329 (2019).

59. Singhal, R., Prata, I.O., Bonnell, V.A. & Llinás, M. Unraveling the complexities of ApiAP2 regulation in Plasmodium falciparum. Trends in Parasitology 40, 987–999 (2024).

60. Elemento, O., Slonim, N. & Tavazoie, S. A universal framework for regulatory element discovery across all genomes and data-types. Molecular cell 28, 337–337 (2007).

61. Gupta, S., Stamatoyannopoulos, J.A., Bailey, T.L. & Noble, W.S. Quantifying similarity between motifs. Genome Biology 8, 1–9 (2007).

62. Wang, J. et al. Haem-activated promiscuous targeting of artemisinin in Plasmodium falciparum. Nat Commun 6, 10111 (2015).

63. Meshnick, S.R. Artemisinin: mechanisms of action, resistance and toxicity. Int J Parasitol 32, 1655–60 (2002).

64. Pires, C.V. et al. Oxidative stress changes the effectiveness of artemisinin in Plasmodium falciparum. mBio 15, e0316923 (2024).

65. Go, K.D. et al. Antimalarial drug artemisinin stabilizes PfRACK1 binding to the ribosome. Structure 33, 1386–1397 e5 (2025).

66. Tripathi, J. et al. The artemisinin-induced dormant stages of Plasmodium falciparum exhibit hallmarks of cellular quiescence/senescence and drug resilience. Nature Communications *2024* 15:*1* 15, 1-17 (2024).

67. Zhang, M. et al. The apicoplast link to fever-survival and artemisinin-resistance in the malaria parasite. Nat Commun 12, 4563 (2021).

68. Ashley, E.A. et al. Spread of Artemisinin Resistance in *Plasmodium falciparum* Malaria. New England Journal of Medicine 371, 411–423 (2014).

69. Mok, S. et al. Drug resistance. Population transcriptomics of human malaria parasites reveals the mechanism of artemisinin resistance. *Science (New York*, N.Y.) 347, 431–435 (2015).

70. Kucharski, M. et al. Short tandem repeat polymorphism in the promoter region of cyclophilin 19B drives its transcriptional upregulation and contributes to drug resistance in the malaria parasite Plasmodium falciparum. PLoS Pathog 19, e1011118 (2023).

71. Han, J., Munro, J.E., Kocoski, A., Barry, A.E. & Bahlo, M. Population-level genome-wide STR discovery and validation for population structure and genetic diversity assessment of Plasmodium species. PLoS Genet 18, e1009604 (2022).

72. Vincent, F. et al. Phenotypic Drug Discovery: Recent successes, lessons learned and new directions. Nature reviews. Drug discovery 21, 899–899 (2022).

73. Gamo, F.J. et al. Thousands of chemical starting points for antimalarial lead identification. Nature *2010* 465:7296 465, 305-310 (2010).

74. Guiguemde, W.A. et al. Global Phenotypic Screening for Antimalarials. Chemistry & Biology 19, 116–129 (2012).

75. Spangenberg, T. et al. The Open Access Malaria Box: A Drug Discovery Catalyst for Neglected Diseases. PLoS ONE 8, e62906–e62906 (2013).

76. Trager, W. On the establishment in culture of isolates of Plasmodium falciparum. Trans R Soc Trop Med Hyg 84, 466 (1990).

77. Lambros, C. & Vanderberg, J.P. Synchronization of Plasmodium falciparum erythrocytic stages in culture. Journal of Parasitology 65, 418–420 (1979).

78. Martin, M. Cutadapt removes adapter sequences from high-throughput sequencing reads. EMBnet.journal 17, 10–12 (2011).

79. Kim, D., Paggi, J.M., Park, C., Bennett, C. & Salzberg, S.L. Graph-based genome alignment and genotyping with HISAT2 and HISAT-genotype. Nature Biotechnology *2019* 37:*8* 37, 907–915 (2019).

80. Li, H. et al. The Sequence Alignment/Map format and SAMtools. Bioinformatics 25, 2078–2079 (2009).

81. Pertea, M. et al. StringTie enables improved reconstruction of a transcriptome from RNA-seq reads. Nature Biotechnology *2015* 33:*3* 33, 290–295 (2015).

82. Benaglia, T., Chauveau, D., Hunter, D.R. & Young, D.S. mixtools: An R Package for Analyzing Mixture Models. Journal of Statistical Software 32, 1–29 (2010).

